# Mast cell regranulation involves a metabolic switch promoted by the interaction between mTORC1 and a glucose-6-phosphate transporter

**DOI:** 10.1101/2021.10.09.463777

**Authors:** Jason A Iskarpatyoti, Jianling Shi, Abhay P S Rathor, Yuxuan Miao, Soman N Abraham

## Abstract

Mast cells (MCs) are highly granulated tissue resident hematopoietic cells and because of their capacity to degranulate and release many proinflammatory mediators, they are major effectors of chronic inflammatory disorders including asthma and urticaria. As MCs have the unique capacity to reform their granules following degranulation *in vitro,* their potential to undergo multiple cycles of degranulation and regranulation *in vivo* has been linked to their pathogenesis. However, it is not known what factors regulate MC regranulation let alone if MC regranulation occurs *in vivo*. Here, we report that IgE-sensitized mice can undergo multiple bouts of regranulation, following repeated anaphylactic reactions. mTORC1, a critical nutrient sensor that activates protein and lipid synthesis, was found necessary for MC regranulation. mTORC1 activity in MCs was regulated by a glucose-6-phosphate transporter, Slc37a2, which was found to be necessary for increased glucose-6-phosphate and ATP levels during regranulation, two upstream signals of mTOR. Slc37a2 is highly expressed at the cell periphery early during regranulation where it appears to colocalize with mTORC1. Additionally, this transporter was found to concentrate extracellular metabolites within endosomes which are trafficked directly into nascent granules. Thus, the metabolic switch associated with MC regranulation is mediated by the interactions of a cellular metabolic sensor and a transporter of extracellular metabolites into MC granules.

## Introduction

Mast cells (MCs) are highly granulated tissue resident cells of hematopoietic lineage, located at the host-environment interface^1,2^. Although a primary physiologic role appears to be promoting innate and adaptive immune response to infectious agents^3–5^, they are best known for promoting chronic inflammatory disorders such as urticaria, anaphylaxis, arthritis, and asthma^6–9^. Much of the pathogenic contribution of MCs is linked to their capacity to release a battery of bioactive mediators that are stored within granules such as lysosomal hydrolases, amines, cytokines, chemokines, proteases and proteoglycans for sustained periods^10^. These mediators, upon release from extracellular MC granules, in a process called degranulation, promote the recruitment of various immune cells from circulation into the tissue^4^. However, when MC release of mediators is excessive or unregulated, significant tissue damage can result.

Since MCs are long lived cells, having been described to survive *in vivo* for several months^11,12^, another MC trait thought to promote its pathogenesis is its unique capacity to regranulate, in place, following degranulation. Indeed, *in vitro* studies have reported that MC degranulation is accompanied by regranulation^13,14^. This suggests that MCs have the potential to undergo multiple cycles of degranulation and regranulation *in vivo* thereby markedly enhancing their capacity to contribute to pathologies found in chronic allergic conditions such as asthma or atopic dermatitis where significant changes to tissues homeostasis are observed^15,16^. However, at this time, it is not known if multiple cycles of MC degranulation and regranulation can occur *in vivo* and if so, what factors regulate this activity.

While the functional relevance of MC degranulation is well established, the biogenesis of MC granules remains less well known. Investigations of MC granule biogenesis have been limited to mostly microscopy-based studies and they have revealed that granule formation is initiated by the formation of uniform-sized progranules within MCs which bud off from the trans-golgi^17,18^. Thereafter, progranules fuse with each other forming larger, immature granules which ultimately associate with endosomes that are positive for CD63, a secretory lysosomal membrane marker^19–21^. Biochemical studies of isolated MC granules have shown them to contain a dense core comprising of negatively charged serglycin proteoglycans which avidly bind to positively charged granule components such as chymase and tryptase^22^. Cumulatively, these multiple ionic interactions stabilize the granule structure and keep them from rapidly disaggregating immediately upon release by MCs allowing distant communications within the body^23,24^.

Conceivably, mechanisms involved in the initial biogenesis of granules, such as protein and lipid synthesis followed by intracellular trafficking of progranules and vesicles, and by the packaging of granule components within MC granules are also involved in MC regranulation. Since the bulk of these activities will require significant energy demands, we hypothesized that following degranulation, marked metabolic reprogramming within MCs would occur to accommodate regranulation. It is well known that mTOR, an evolutionarily conserved ser/thr kinase, is a critical regulator of the metabolic switch that occurs during T cell differentiation^25,26^. Specifically, mTORC1, one of two complexes formed by mTOR, is activated upon sensing sufficient glucose and amino acid levels which leads to downstream phosphorylation of 4E-BP and S6K which in turn leads to protein and lipid synthesis^27^. Additionally, mTORC1 activation is required for increased glycolysis in T cells during differentiation^28^. Based on these observations, we propose that mTORC1 activation is necessary for MC regranulation. Here, we sought to validate this notion and as well as identify additional factors in MCs that might contribute to promoting MC regranulation.

## Methods

### Mice and anaphylaxis experiments

Six- to eight-week-old female C57BL/6J and Raptor^f/f^ mice were purchased from The Jackson Laboratory and housed in Duke University animal facilities. Raptor^f/f^-ER Cre mice were generously donated by Dr. Xiao Ping Zhong at Duke University. We used six- to twelve-week-old C57BL/6J and Raptor^f/f^ ER-Cre mice, as well as Cre negative littermate controls for our experiments. Raptor ER-Cre mice and littermate controls were treated with 2 mg tamoxifen (Tocris) in 100 μL corn oil via gavage on 5 consecutive days. All animal experiments were performed according to approved protocols by the Duke University Institute Animal Care and Use Committee.

For anaphylaxis experiments, mice were sensitized i.p. with 10 μg anti-TNP IgE (BD Biosciences) in 100 μL sterile PBS. Mice were Ag-challenged 16 hrs later i.p. with 50 μg TNP-OVA (Protein Sciences) in 100 μL sterile PBS. Control mice were challenged with sterile PBS. Core body temperatures were monitored using a rectal thermometer immediately prior to Ag-challenge and then every 15 minutes for up to 3 hrs. Repeat Ag-challenges were separated by 7 days and mice were resensitized with 10 μg anti-TNP IgE and Ag-challenged 16 hrs later with 50 μg TNP-OVA.

### Cell lines

BMMCs were derived from femur bone marrow of WT C57BL/6J, Raptor^f/f^-ER Cre, and littermate controls negative for Cre. Cells were cultured in RPMI-1640 cell media containing 2 mM L-glutamine (ThermoFisher), 10% heat-inactivated fetal bovine serum (FBS, Hyclone), 1 mM nonessential amino acids (ThermoFisher), 25 mM HEPES (ThermoFisher), 1 mM sodium pyruvate (ThermoFisher), 1X Antibiotic-Antimycotic (ThermoFisher), 5 ng/mL recombinant murine IL-3 (R&D Systems, and 5% stem cell factor-containing supernatant from CHO-KL cells for 4-6 weeks until maturation. CHO-KL cells were a gift from M. Arock (Laboratoire de Biologie et Pharmacologie Appliquée, Paris, France). RBL-2H3 cells (ATCC) were cultured in MEM media containing Earl salt’s, 2 mM L-glutamine (ThermoFisher), 15% heat-inactivated FBS, and 1 mM sodium pyruvate.

Generation of RBL-2H3 cells stably expressing serglycin-mCherry was described previously^29^. To generate Slc37a2-EGFP expressing RBLs, Slc37a2 was amplified by PCR from BMMCs cDNA and inserted into a plEGFP-C1 plasmid (Takara Bio). Slc37a2 plEGFP-C1 plasmid or plasmid vector control were transfected into Amphopack-293 cells (BD Biosciences) to produce viral particles that were used to infect RBL-2H3 cells or RBL-2H3 cells stably expressing serglycin-mCherry. Slc37a2-EGFP stably expressing cells were selected using neomycin.

### Flow cytometry analysis

Mice were Ag-challenged one, two, or three times, as previously described. 24 hrs after the final Ag-challenge, peritoneal cells were collected after lavage with 5 mL cold 10 mM EDTA in PBS. Samples were washed with PBS containing 5 mM EDTA and 3% FBS (FACS Buffer) and then blocked with 1% anti-mouse CD16/CD32 (BD biosciences), 5% normal rat serum and 5% normal mouse serum in FACS buffer for 10 minutes at RT. Then, surface staining was performed using 7-AAD (Biolegend) and fluorochrome-tagged antibodies to CD45 and CD117 (both from Biolegend), and CD11b, Ly6G, SiglecF, Ly6C, and F4/80 (all from BD Biosciences). Intracellular staining for Avidin (BD Biolegend) was performed using the BD Cytofix/Cytoperm Fixation/Permeabilization Kit and following manufacturer’s instructions. Data was collected using BD Fortessa X-20 (BD Biosciences) and analyzed with FlowJo software (TreeStar). The gating strategy used is described in Supplemental Figure 1.

### Immunofluorescence staining and microscopy

For visualization of BMMCs and peritoneal cells, cells were cytospun onto charged glass slides. For RBL-2H3 cells, cells were seeded onto sterilized glass coverslips and allowed to adhere overnight. Cells were then fixed using 4% paraformaldehyde for 20 minutes at room temperature (RT). Cells were washed in PBS three times and permeabilized in 0.1% saponin (Sigma) and blocked in 1% BSA (USBiological Life Sciences) in PBS for 20 minutes at RT. Cells were then incubated with 1% anti-CD63 (MBL), 0.5% anti-Slc37a2 (GeneTex), 1% anti-serotonin (Dako), or 1% anti-mTOR (EMD Millipore) overnight at 4°C. The following day, cells were washed in 0.1% saponin and 1% BSA in PBS three times and incubated with 0.25% fluorescent secondary antibodies (Jackson ImmunoResearch) or 0.2% avidin conjugated with FITC (BD Biosciences) for 1 hr at RT. Cells were then washed twice with 0.1% saponin and 1% BSA in PBS and twice with PBS. Slides were mounted with ProLong Diamond (Invitrogen). Images were acquired using the Leica Thunder Imager and analyzed by ImageJ.

### β-hexosaminidase assay

For BMMC degranulation assays, BMMCs were plated at 1×10^5^ cells/well in a 96 well V-bottom plate. Cells were washed in Tyrode’s buffer (135 mM NaCl, 5 mM KCl, 1.3 mM CaCl_2_, 1 mM MgCl_2_, 5.5 mM glucose, 1% BSA, 20 mM HEPES, pH 7.4) and then incubated with 1 μg/mL ionomycin (Sigma) in Tyrode’s buffer for 1 hr at 37 °C. BMMCs in Tyrode’s buffer alone were used as a negative control. After incubation, supernatants were collected and cell pellet was lysed in 0.1% Triton-100X (Spectrum Chemical) in Tyrode’s buffer. Collected supernatant and lysed pellet were separately incubated with 3.4 mg/mL p-nitrophenyl-N-acetyl-β-D-glucosaminide (NAG, Sigma) in citrate buffer (pH 4.5) for 1 hr at 37°C. Reactions were stopped at 1 hr with 0.1 M carbonate buffer (pH 10). Colorimetric measurements were performed at 405 nm using a Synergy H1 microplate reader (BioTek). Percent degranulation was determined as percent β-hexosaminidase activity in supernatant compared to total β-hexosaminidase activity in supernatant and pellet.

To measure regranulation, 1 hr post-activation with ionomycin, BMMCs were resuspended in BMMC media and incubated at 37°C for 48 hrs, unless otherwise noted. Total β-hexosaminidase activity was measured by plating 1×10^5^ cells/well in a 96 well V-bottom plate. BMMCs were washed in Tyrode’s buffer then lysed in 0.1% Triton-100X (Spectrum Chemical) in Tyrode’s buffer. Total β-hexosaminidase activity in lysed pellet was determined as described above.

For regranulation experiments using 1 mM 2-deoxyglucose (Sigma), 33 μM nocodazole (Sigma), or 20 nM torin (Tocris), BMMCs were treated 1 hr post degranulation with each respective inhibitor in BMMC medium. Total β-hexosaminidase activity was measured 48 hrs post degranulation.

### Western Blot

Cells were lysed at indicated time points post degranulation using RIPA buffer (Sigma) supplemented with protease inhibitor cocktail (Sigma) and phosphatase inhibitor cocktails 2 and 3 (Sigma). Lysates were incubated at 4°C for 30 minutes and supernatants were collected after centrifugation at 20,000xG at 4°C for 15 minutes. Total protein concentrations were determined using the Pierce™ Coomassie (Bradford) Protein Assay Kit according to manufacturer’s instructions (ThermoFisher). Equal concentrations of cell lysates were prepared in Laemmli buffer (BioRad) containing a 5% β-mercaptoethanol (Sigma) and denatured at 95°C for 5 minutes. Proteins were separated on a 4% to 20% mini-PROTEAN TGX precast gel (Biorad) and transferred onto a nitrocellulose membrane by wet transfer. Membranes were blocked in 5% BSA or 5% milk dissolved in TBS (Biorad) containing 0.1% Tween 20 (Sigma) for 45 minutes and then incubated at 4°C overnight with primary antibodies in blocking buffer. Membranes were washed five times in TBST then incubated with secondary antibody conjugated to horseradish peroxidase (HRP) for 1 hr at RT. Membranes were washed five times in TBST then proteins were detected using enhanced chemiluminescent HRP substrate (ThermoFisher). Antibody dilutions were 1:1000 for anti-mTOR, anti-p-mTOR S2448, anti-S6K, and anti-p-S6K S371 (Cell Signaling), and 1:5000 for anti-β-actin (Sigma).

### Toluidine Blue Assay

72 hrs after mice were Ag-challenged, peritoneal cells were collected after lavage with 5 mL cold 10 mM EDTA in PBS. 100 μLs of cells were cytospun onto glass slides and fixed with Carnoy’s fixative (60% ethanol, 30 % chloroform, and 10% glacial acetic acid) for 20 minutes at RT and stained with 0.1% toluidine blue dissolved in PBS containing 0.5N HCl for 20 min. Granulated MCs were visualized under a brightfield microscope.

### Microarray

RNA was collected from untreated BMMCs or BMMCs 4 hrs after treatment with 1 μg/mL ionomycin using the RNeasy Mini Kit. Total RNA was assessed for quality with Agilent 2100 Bioanalyzer G2939A (Agilent Technologies) and Nanodrop 8000 spectrophotometer (Thermo Scientific/Nanodrop). Hybridization targets were prepared with the Affymetrix Whole Transcriptome Plus Kit (Affymetrix) and Affymetrix GeneChip Whole Transcriptome Terminal Labeling Kit (included with Affy Whole Transcriptome Plus) from total RNA, hybridized to GeneChip® Mouse Transcriptome Array 1.0 in Affymetrix GeneChip® hybridization oven 645, washed in Affymetrix GeneChip® Fluidics Station 450 and scanned with Affymetrix GeneChip® Scanner 7G according to standard Affymetrix GeneChip® Hybridization, Wash, and Stain protocols. (Affymetrix, Santa Clara, CA).

Affymetrix Gene Chip microarray data was processed using the *oligo*^*30*^ package in the Bioconductor suite^31^ from the R statistical programming environment. Log-scale Robust Multiarray Analysis (RMA) was used to normalize the data and eliminate systematic differences across the arrays. Differential expression between the naïve and the wild-type samples was calculated with an empirical Bayes moderated test statistic using the *limma* package^32^. The False Discovery Rate (FDR) was used to control for multiple hypothesis testing. Gene set enrichment analysis^33^ (GSEA) was performed using the *fgsea*^*34*^ package to identify gene ontology terms and pathways associated with the differentially expressed genes.

### RT-qPCR

Total RNA was extracted from BMMCs at indicated time points post ionomycin treatment using the RNeasy mini kit (Qiagen) according to manufacturer’s instructions. cDNA was then synthesized using the iScript cDNA synthesis kit (BioRad) and following manufacturer’s instruction. Finally, RT-qPCR was performed using iQ SYBR Green Supermix (BioRad) and run on in triplicate on a StepOnePlus Real-Time PCR System (Applied Biosciences) and analyzed by StepOne Software v2.3. mRNA amounts were normalized using β-actin and calculated using the 2^−ΔΔCT^ method. The following primer pairs purchased from IDT were used: Slc37a2, 5’-GCTGCTCCCATGATGTTCCT-3’ and 5’-TTGGCGTTACCCTTCAGGCT-3’, β-ACTIN, 5’-GATTACTGCTCTGGCTCCTAGC-3’ and 5’-GACTCATCGTACTCCTGCTTGC-3’.

### siRNA Treatment

Slc37a2 was knocked down in BMMCs using siGENOME SMARTpool siRNAs (Dharmacon). Transfection of siRNAs into BMMCs was performed using the Amaxa Nucleofector Kit T (Lonza). BMMCs were pelleted and resuspended at a concentration of 2×10^6^ cells/100 μL Nucleofector T solution. 300 nM Slc37a2 siRNAs or Silencer™ negative control siRNA (Ambion) was added to the solution. BMMCs were nucleofected using the Amaxa Nucleofector Program U-023. Transfected BMMCs were then resuspended in cell media for 48 hrs at 37°C prior to assaying.

### Dextran Uptake Assay

RBL-2H3 cells stably expressing Slc37a2-EGFP or EGFP vector control seeded onto coverslips, or BMMCs treated with Slc37a2 siRNAs or negative control siRNAs were incubated with 0.5 mg/mL Dextran Texas Red 10,000 MW in cell media for 16 hrs at 37°C. Cells were washed twice with cell media and then incubated an additional 2 hrs in cell media at 37°C. For RBL cells, cells were fixed with 4% paraformaldehyde, washed three times in PBS, and mounted onto a glass slide using Prolong Diamond. For BMMCs, cells were cytospun onto charged glass slides, fixed with 4% paraformaldehyde, washed three times in PBS and then a coverslip was mounted using ProLong Diamond. Images were acquired using the Leica Thunder Imager and analyzed by ImageJ.

### Glucose-6-Phosphate Assay

BMMCs, treated with Slc37a2 siRNA or control siRNA, were activated using 1 μg/mL ionomycin in Tyrode’s buffer. 1 hr after ionomycin treatment, BMMCs were washed three times and resuspended in BMMC medium. G6P concentrations were determined at 2 hrs and 6 hrs post activation and compared to unactivated controls using a G6P kit (Sigma) and following manufacturer’s instructions.

### Seahorse XF Assays

For Slc37a2 siRNA knockdown experiments, Slc37a2 siRNA or control siRNA treated BMMCs were sensitized overnight with 1 μg/mL anti-TNP IgE. Cells were washed three times and resuspended in Tyrode’s buffer containing 10 ng/mL TNP-OVA or PBS. After 1 hr, cells were washed three times and resuspended in BMMC media. For nocodazole experiments, BMMCs were treated with 1 μg/mL ionomycin or PBS in Tyrode’s buffer. 1 hr after ionomycin treatment, BMMCs were washed three times and resuspended in BMMC medium containing 33 μM nocodazole. For BMMC adherence, XF96 cell culture microplates were precoated with Cell-Tak following manufacturer’s instruction. At 6 hrs post TNP-OVA or ionomycin treatment, BMMCs were seeded onto Cell-Tak treated XF96 cell culture microplates at a density of 1×10^5^ cells/well. Basal oxygen consumption rates (OCR) and extracellular acidification rates (ECAR) were measured using the Seahorse XF96e Analyzer followed by sequential addition of 1 μM oligomycin (Sigma), 0.5 μM carbonyl cyanide 4-(trifluoromethoxy) phenylhydrazone (FCCP, Sigma), and 0.75 μM rotenone (Sigma) with 1.5 μM antimycin A (Sigma). Basal respiration, basal glycolysis, ATP production, and maximal respiration were evaluated based on the manufacturer’s instructions.

### Statistics

Statistical analyses were performed using GraphPad Prism v.9.0.0 (GraphPad Software). For comparison between two groups, results were analyzed using a two-tailed unpaired *t* test. For comparison between multiple groups, results were analyzed using a two-tailed ANOVA with Tukey’s post test. A *p*-value less than 0.05 was considered statistically significant.

## Results

### MCs can undergo multiple bouts of degranulation and regranulation in mice

To date, the capacity of MCs to undergo more than one round of degranulation *in vivo* has not been studied. Here, we investigated if IgE-sensitized peritoneal MCs can undergo multiple rounds of degranulation *in vivo* following serial exposure to antigen (Ag). A mouse model of anaphylaxis was used to assess the magnitude of the inflammatory response, which is directly induced by and correlated to the degranulation of IgE sensitized MCs^29,35^. To induce MC degranulation, we passively sensitized mice with 10 μg TNP-specific IgE i.p. then, 16 hours later, challenged these mice with 50 μg TNP-OVA Ag i.p. All of the sensitized mice experienced anaphylaxis as shown by a drastic drop in core body temperature (Fig. 1a, 1^st^ Ag challenge). We observed that temperatures begin to drop in the mice within 15 minutes of antigen challenge. Temperatures continued to drop for 30 minutes and thereafter the mice began to recover and return to baseline core body temperature (Fig. 1a).

**Figure 1.**
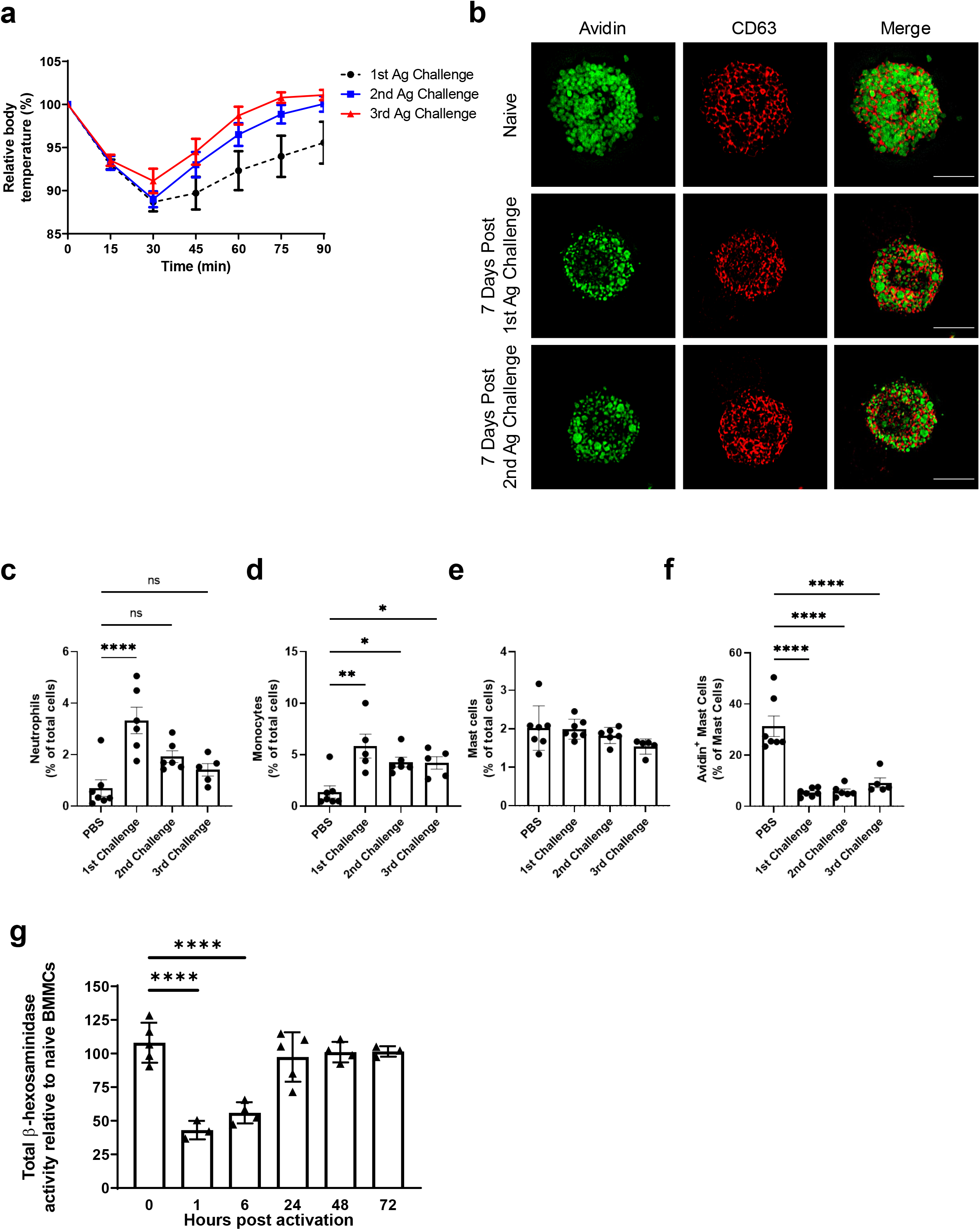
Regranulated MCs are capable of causing repeat anaphylaxis *in vivo*. (**a**) Mice were sensitized with 10 μg TNP-specific IgE i.p. and challenged 16 hrs later with 50 μg TNP-OVA i.p. Mice were then sensitized and Ag challenged an additional two more times, waiting a week between challenges. Core body temperatures were measured after each challenge using a rectal thermometer and expressed as relative body temperature compared to initial temperatures (*n* ≥ 7 mice). (**b**) Peritoneal lavages were obtained from naive mice or 7 days after mice were Ag challenged a first or second time. Cells were cytospun onto coverslips and analyzed by immunofluorescence with staining for Avidin (green) and CD63 (red). Images are representative of at least 2 independent experiments. Scale bars represent 10 μm. (**c-f**) Peritoneal lavages were obtained 24 hrs after the first, second, or third Ag challenge or from PBS treated control mice and analyzed by flow cytometry. Quantification of these results is shown for (**c**) CD45^+^CD11b^+^Ly6G^+^ neutrophils (**d**) CD45^+^CD11b^+^Ly6C^+^ monocytes, (**e**) CD45^+^cKit^+^ mast cells, and (**f**) CD45^+^cKit^+^Avidin^Hi^ intracellular mast cell staining. (*n* ≥ 6 mice) (**g**) BMMCs were activated using 1 μg/mL ionomycin in Tyrode’s buffer for 1 hr. Cells were washed three times and resuspended in cell medium. Total β-hexosaminidase was measured at 1, 6, 24, 48, and 72 hrs post activation and compared to untreated BMMC controls. Data are representative of at least 3 independent experiments. Data were analyzed by a 1-way ANOVA with a Tukey’s post-test (**b-e**), or a 1-way ANOVA with a Dunnett’s post-test comparing each group with untreated BMMC controls. \**P* < 0.05; \*\**P* < 0.01; \*\*\**P* < 0.001; \*\*\*\**P* < 0.0001; ns, not significant.

Since MCs are long-lived cells that have demonstrated the capacity to regranulate *in vitro*^*13,14*^, we reasoned that recently degranulated mouse peritoneal MCs should regranulate *in vivo* and participate in additional anaphylactic reactions when challenged a second time and even a third time with Ag. To investigate if recently degranulated MCs could regranulate *in vivo,* we collected peritoneal MCs from mice that had experienced anaphylaxis and naïve mice and examined them via microscopy for their state of granulation. We selected day 7 post Ag-challenge as we had previously established that by day 7, all of the inflammatory reactions induced by the initial challenge experiment had subsided in the peritoneum (data not shown). MCs from both groups of mice were stained with anti-CD63 antibody, a probe for MC granule membranes, and Avidin-FITC a probe for the MC granule component, heparin^36^. In naïve mice, Avidin-FITC^+^ and CD63^+^ MC granules occupied most of the cytoplasm of peritoneal MCs and were of relatively uniform size (Fig. 1b, top row). Although the level of granulation in MCs from Ag-challenged mice 7 days prior were comparable to naïve MCs, the granule size and shape was far more heterogeneous (Fig. 1b, middle row). As we would expect newly recruited MCs to appear similar to naïve MCs, these results confirm that 7 days after Ag-challenge, MCs are regranulated *in vivo* although the granules were not uniform in size.

Next, we sought to examine if these regranulated MCs could function regularly by mediating anaphylaxis. A week after the initial sensitization and Ag challenge, we passively sensitized mice again with 10 μg TNP-specific IgE i.p. and then challenged these mice with 50 μg TNP-OVA i.p. We found that mice challenged 7 days previously exhibited a comparable drop in core body temperature, reaching the lowest core body temperatures by 30 minutes (Fig.1a, 2^nd^ Ag challenge). 7 days after the second challenge, peritoneal MCs were examined and we found that similar to MCs seven days after the initial Ag challenge, MC granule size and shape were far more heterogeneous than observed in naïve mice (Fig. 1b, bottom row). Furthermore, a similar drop in body temperature was observed when these mice were sensitized and challenged for a third consecutive time with Ag 7 days later (Fig. 1a, 3^rd^ Ag challenge). Interestingly, we observed a trend towards faster recovery time from anaphylaxis after each Ag-challenge. These observations reveal for the first time that MCs can undergo repeated bouts of degranulation without significant loss in degranulation (or regranulation) capacity.

To further confirm that MCs can undergo multiple bouts of degranulation and regranulation without significant loss in functional capacity, we compared immune cell recruitment into the mouse peritoneal cavity, the site of MC activation, after each Ag challenge. Mice were euthanized 24 hours after the first, second or third Ag-challenge and peritoneal cells were collected by lavage. Inflammatory response was determined by flow cytometry (Supplemental Fig. 1). We found that 24 hours after the initial Ag-challenge, the percentage of CD45^+^CD11b^+^Ly6G^+^ neutrophils, and CD45^+^Ly6G^−^Ly6C^+^ monocytes were increased compared to PBS treated controls (Fig. 1c and d). Notably, the second and third Ag-challenge, while causing anaphylaxis, had similar percentage of neutrophils in the peritoneum compared to PBS-control mice while maintaining increased monocyte recruitment. Importantly, the percentage of cKit^+^ MCs in the peritoneum remained similar after each Ag-challenge compared to PBS controls (Fig. 1e). As expected, intracellular staining with avidin, a MC-specific granule marker, showed that after each Ag-challenge there were significantly fewer Avidin^Hi^ MCs compared to PBS control mice, suggesting that 24 hours after each Ag challenge, MCs were mostly degranulated (Fig. 1f). Taken together, this indicates that the second and third Ag-challenge resulted in MC degranulation capable of producing robust anaphylaxis inflammatory response.

These findings reveal, for the first time, that MCs have the innate capacity to regranulate *in vivo* after multiple bouts of degranulation, a distinct property that can potentially contribute to sustaining chronic MC mediated inflammatory reactions.

### Kinetics of MC regranulation *in vitro*

Having established that MC regranulation can occur *in vitro* and *in vivo*, we sought to elucidate the molecular basis of this distinct activity. First, we investigated the kinetics of MC regranulation. For these studies, we employed cultured primary bone marrow-derived MCs (BMMCs) from mice. To ensure that our studies of MC regranulation were broadly applicable to other modes of MC degranulation we induced MCs degranulation employing ionomycin, a Ca^2+^ ionophore that induces calcium flux a critical signaling event preceding MC degranulation regardless of the agonist employed. We assessed degranulation and regranulation by measuring residual cellular content of β-hexosaminidase, a component of the granule that is released upon degranulation and is resynthesized within granules when MCs regranulate. As expected, following degranulation, total β-hexosaminidase levels in BMMCs were significantly reduced compared to untreated controls (Fig. 1g). Six hours post degranulation, β-hexosaminidase levels remained largely unchanged, however, between 24 and 48 hours post degranulation, BMMCs had recovered β-hexosaminidase levels that were comparable to levels seen prior to degranulation. Thus, *in vitro* MC regranulation takes between 24 and 48 hours to complete.

### mTORC1 is necessary for MC regranulation

For MCs to fully regranulate, significant protein and lipid synthesis must occur followed by vesicle trafficking, and proper packaging of various granule components into granule structures. These processes exert increased metabolic demands on the cell, requiring substantial metabolic reprogramming to increase available energy. In this regard, a potentially important cellular regulator of metabolic reprogramming is mTOR, a kinase that has previously been shown to act as a metabolic switch in T cell activation as well as several innate immune cell functions^25,26,37^. To investigate a possible role for mTOR in MC regranulation, we induced MC degranulation employing ionomycin as before and measured mTOR activity by Western blot. We observed that at 2 hours post degranulation, mTOR had increased phosphorylation (Fig. 2a). Furthermore, S6K a serine/threonine kinase, and a downstream target of mTORC1, also appeared to be phosphorylated at the same time point (Fig. 2a). To assess the functional role of mTOR during regranulation, BMMCs were treated with 20 nM torin, a selective inhibitor of mTOR activation^38^, prior to degranulation with ionomycin. Torin-pretreated BMMCs exhibited no significant difference in total percent release of β-hexosaminidase upon degranulation (Fig. 2b). However, BMMCs maintained in 20 nM torin following degranulation were inhibited in their capacity to regranulate based on the limited amounts of total β-hexosaminidase found in BMMCs 48 hours post degranulation (Fig. 2b) suggesting that although mTORC1 plays no significant role in MC degranulation, its activation is necessary for MC regranulation.

**Figure 2.**
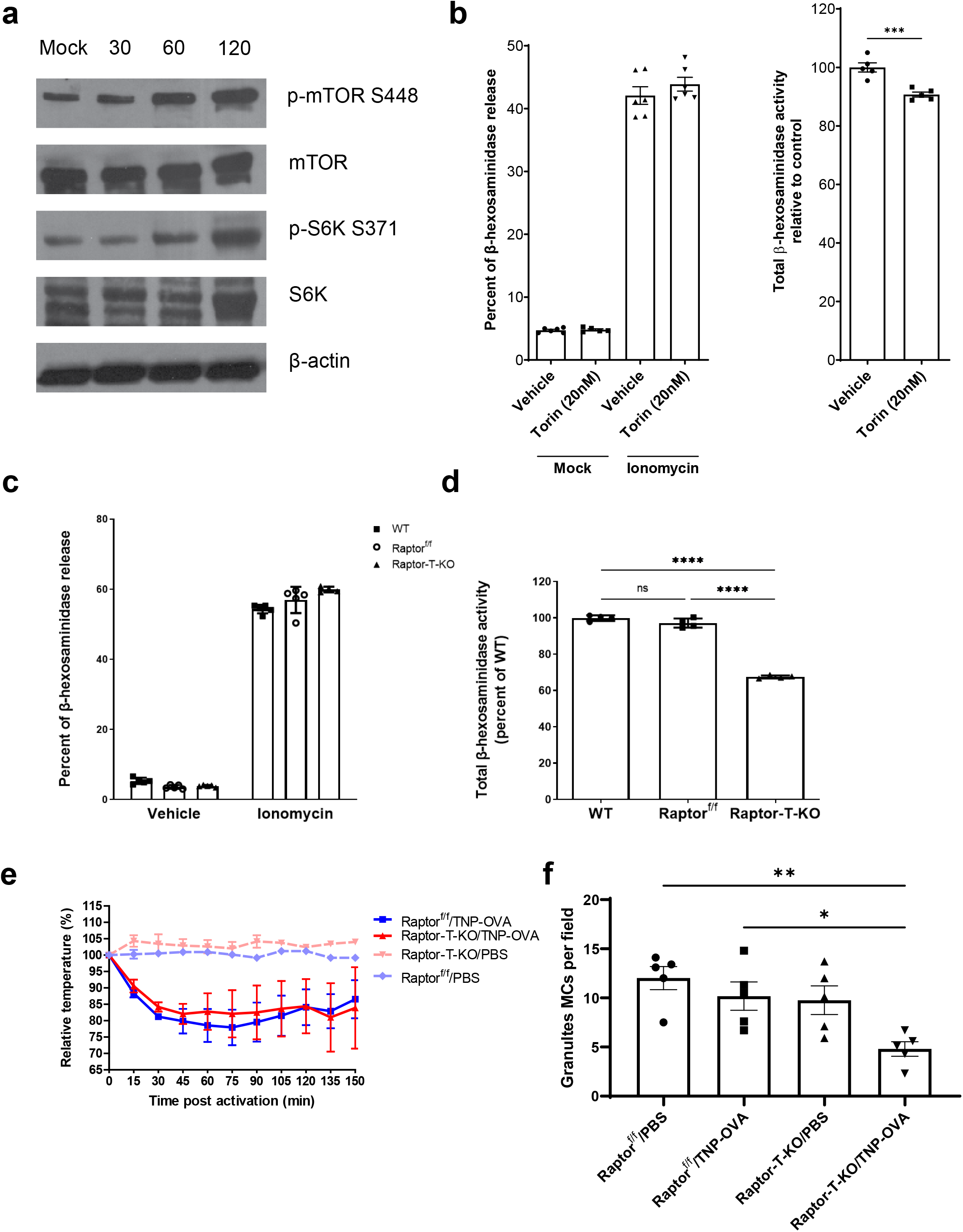
mTORC1 is necessary for MC regranulation *in vitro* and *in vivo*. (**a**) Western blot analysis of untreated BMMCs and BMMCs 30 min, 60 min, or 120 min after activation by 1 μg/mL ionomycin. Immunoblots were incubated overnight with anti-p-mTOR S2448, anti-mTOR, anti-p-S6K S371, anti-S6K, and anti-β-actin. Two independent experiments were performed with similar results. (**b, left**) BMMCs were pretreated with 20 nM torin or DMSO (vehicle control) and then activated using 1 μg/mL ionomycin in Tyrode’s buffer or treated with Tyrode’s buffer alone (Mock) for 1 hr. β-hexosaminidase release was measured and expressed as a percent of total. (**b, right**) BMMCs were activated using 1 μg/mL ionomycin in Tyrode’s buffer then cells were washed three times and resuspended in cell medium containing 20 nM torin or DMSO. Total β-hexosaminidase was measured at 48 hrs post ionomycin treatment. Results are displayed as a percent of vehicle control. (**c**) BMMCs from C57BL/6J mice (WT), Raptor^f/f^, or Raptor-T-KO mice were activated using 1 μg/mL ionomycin in Tyrode’s buffer or Tyrode’s buffer alone (vehicle control) for 1 hr. β-hexosaminidase release was measured and expressed as a percent of total. (**d**) BMMCs from C57BL/6J (WT), Raptor^f/f^, or Raptor-T-KO mice were treated with 1 μg/mL ionomycin in Tyrode’s buffer for 1 hr and then washed three times and resuspended in cell medium. Total β-hexosaminidase was measured 48 hrs post activation. Results were normalized to WT BMMCs. Data are representative of three independent experiments. (**d**) Raptor^f/f^ or Raptor-T-KO mice were sensitized with 10 μg TNP specific IgE i.p. and challenged 16 hrs later with 100 μg TNP-OVA i.p. Core body temperatures were measured after each challenge using a rectal thermometer and expressed as relative body temperature compared to initial temperatures. (*n* = 5). (**e**) C57BL/6J (WT), Raptor^f/f^, and Raptor-T-KO mice were sensitized with 10 μg TNP specific IgE i.p. and challenged 16 hrs later with 100 μg TNP-OVA i.p. 72 hrs post Ag challenge, peritoneal lavages were obtained. Cells were cytospun onto coverslips and stained using Toluidine blue. Granules MCs were counted in ten fields of focus per mouse. Error bars represent SEM from *n* = 5 mice. Data were analyzed by a Student’s *t* test (**b**) or a 1-way ANOVA with a Tukey’s post-test (**c-f**). \**P* < 0.05; \*\**P* < 0.01; \*\*\**P* < 0.001.

To extend these findings, we investigated tamoxifen-inducible Raptor knockout (Raptor^f/f^ ER-Cre (Raptor-T-KO)) mice. Raptor (regulatory-associated protein of mammalian target of rapamycin (mTOR)) is a necessary component for the formation of mTORC1. We hypothesized that we could demonstrate the role of mTORC1 by selectively inhibiting MC regranulation without inhibiting the initial granule biogenesis by treating BMMCs derived from these mice with tamoxifen only after granule maturation has been completed. Mature BMMCs from Raptor-T-KO mice were treated with 4-hydroxytamoxifen to knockout Raptor (Supplemental Fig. 2a). Raptor knockout in these BMMCs was confirmed by Western blot (Supplemental Fig 2b.) These BMMCs were then degranulated using ionomycin. We found that tamoxifen-treated Raptor-T-KO BMMCs contained similar amount of β-hexosaminidase compared to littermate control (Raptor^f/f^) BMMCs, which lack the Cre gene. Additionally, Raptor-T-KO and Raptor^f/f^ BMMCs exhibited similar amount of β-hexosaminidase release compared to WT C57BL/6J BMMCs (Fig. 2c). Importantly, 48 hours after degranulation, Raptor-T-KO BMMCs contained significantly reduced amount of β-hexosaminidase compared to Raptor^f/f^ and WT BMMCs confirming that mTORC1 is necessary for MC regranulation but not for degranulation (Fig. 2d).

We next sought to replicate these findings *in vivo*. Raptor-T-KO mice were treated with tamoxifen via gavage once a day for 5 days and knockout was confirmed by PCR (Supplemental Fig. 2c). We then passively sensitized Raptor-T-KO mice and littermate controls i.p. with anti-TNP IgE and Ag-challenged these mice 16 hours later with TNP-OVA i.p. We found that in these mice there was no significant difference in core body temperature drop between Raptor-T-KO mice and littermate controls, suggesting that Raptor-T-KO mice maintain the capacity to undergo anaphylaxis and MCs maintain the capacity to degranulate *in vivo* (Fig. 2e). 72 hours after antigen challenge, peritoneal cells were collected, and granulated MCs were counted by toluidine blue staining which marks granulated MCs. We found that PBS-treated Raptor-T-KO mice had similar levels of granulated MCs compared to littermate controls (Fig. 2f). Notably, Raptor-T-KO mice that were Ag-challenged had a significantly reduced number of granulated MCs compared to littermate controls. These results reveal that in Raptor-T-KO mice, the capacity of MCs to undergo regranulation is impaired. Taken together, our findings confirm that the well-known nutrient sensor mTORC1 is necessary for both *in vitro* and *in vivo* MC regranulation.

### Slc37a2 is upregulated and necessary for MC regranulation

Having identified mTORC1 as a vital regulator of MC regranulation, we sought to identify the basis of this regulation. Our strategy involved using an unbiased approach looking for highly upregulated metabolic genes following MC degranulation and seeing if they interacted with mTORC1. We examined RNA expression in BMMCs 4 hrs after ionomycin induced degranulation. We picked this time-point to selectively identify genes involved in early metabolic activity relating to regranulation while avoiding genes already activated during BMMC degranulation. From our microarray analysis, we identified 44 genes that were differentially regulated in ionomycin treated BMMCs compared to naïve BMMCs, with an adjusted *p*-value less than 0.05 and at least a two-fold change (Fig. 3a). Among those genes, Slc37a2 stood out as a gene involved in glucose metabolism, a process regulated by mTOR activation (Figure 3a). This gene is part of the Slc37 family of genes consisting of Slc37a1, Slc37a2, Slc37a3, and Slc37a4 all of which exhibit sequence homology to the bacterial organophosphate/phosphate (Pi) antiporter^39^. Slc37a2 has been shown to have the capacity to transport glucose-6-phosphate (G6P) across membranes^40^, and is involved in regulating glycolysis in macrophages^41^.

**Figure 3.**
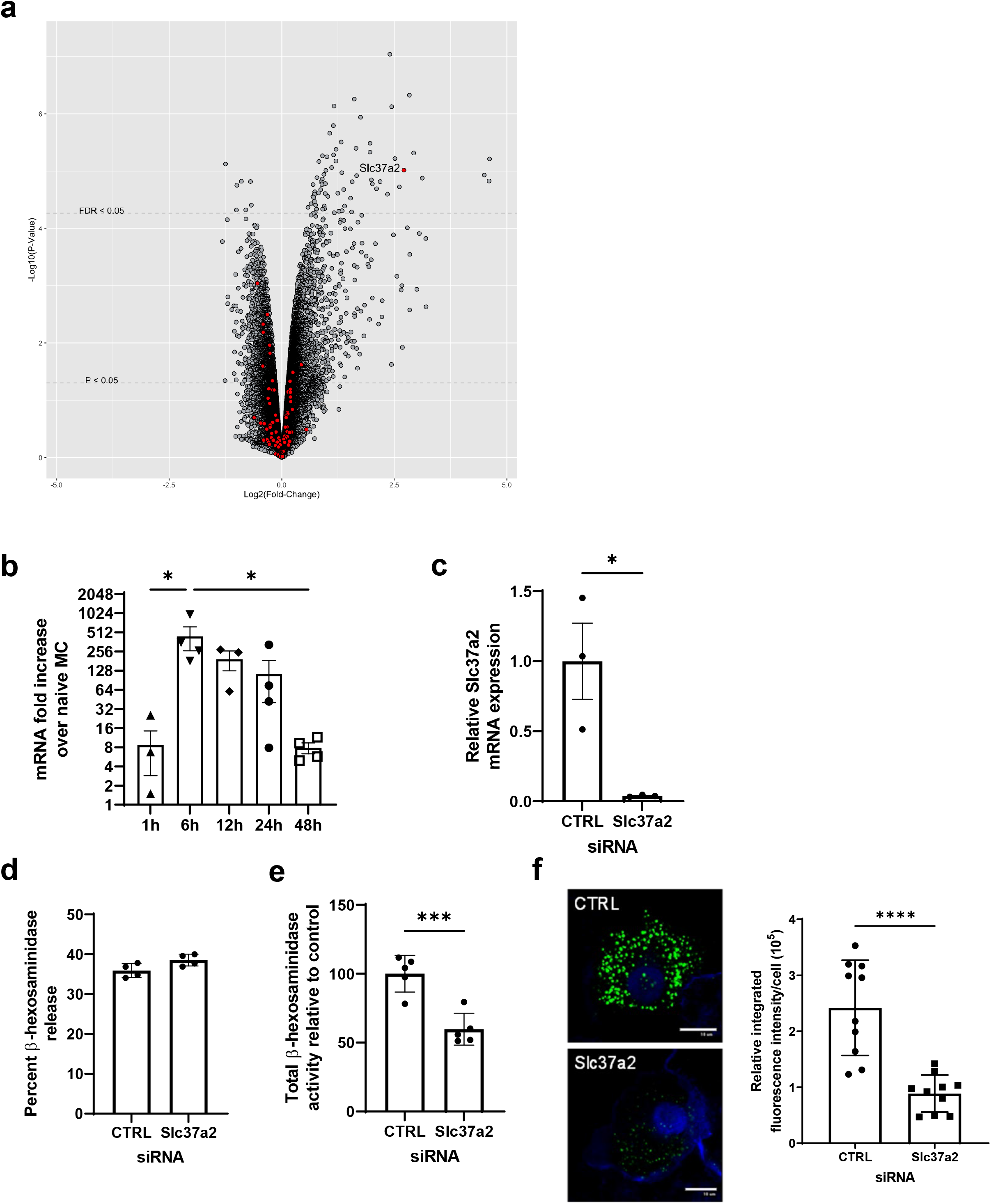
Slc37a2 is a necessary gene in MC regranulation. (**a**) Total RNA was collected from naïve BMMCs and BMMCs 4 hrs after treatment with 1 μg/mL ionomycin. Microarray analysis was performed on RNA samples using three independent samples of RNA for each treatment. Results are displayed in a volcano plot. Genes with a GO Term of Glucose Metabolism Pathway are highlighted in red. (**b**) Slc37a2 mRNA expression was assessed in BMMCs at 1, 6, 12, 24, and 48 hrs post treatment with 1 μg/mL ionomycin by RT-qPCR and normalized to untreated BMMCs. Error bars represent SEM from at least three independent experiments. (**c-f**) BMMCs were treated with Slc37a2 siRNAs or control siRNAs. 48 hrs after treatment, (**c**) Slc37a2 mRNA expression was assessed in BMMCs by RT-qPCR. Results were expressed relative to control siRNA treated BMMCs. (**d**) β-hexosaminidase release was measured after treatment with 1 μg/mL ionomycin in Tyrode’s buffer for 1 hr. Results were expressed as a percent of total. (**e**) BMMCs were treated with 1 μg/mL ionomycin in Tyrode’s buffer for 1 hr and then washed three times and resuspended in cell medium. Total β-hexosaminidase was measured 48 hrs post activation. Results were normalized to control siRNA treated BMMCs 48 hrs after degranulation. (**f**) BMMCs were treated with 1 μg/mL ionomycin in Tyrode’s buffer for 1 hr and then washed three times and resuspended in cell medium. 48 hrs post activation, cells were cytospun onto coverslips followed by fixation and staining for serotonin (green) and phalloidin (blue). Scale bars: 10 μm. Serotonin was quantified using ImageJ; (*n* = 10). Data were analyzed by a 1-way ANOVA with a Tukey’s post-test (**b**), or a Student’s *t* test (**c-f**). \**P* < 0.05; \*\**P* < 0.01; \*\*\**P* < 0.001; \*\*\*\**P* < 0.0001.

We next performed RT-qPCR mRNA analysis of Slc37a2 expression over a 48-hour time course following degranulation and revealed expression peaks at 6 hours post-degranulation, displaying a >400-fold increase in mRNA. Thereafter, expression begins to subside, decreasing significantly and approaching baseline by 48 hours (Fig. 3b).

To determine if Slc37a2 contributes to MC regranulation, we employed siRNAs to knock down its expression in BMMCs and examined the impact of its limited presence on MC regranulation. As shown in Figure 3c, Slc37a2 mRNA levels in BMMCs was markedly reduced compared to control siRNA treated BMMCs. Furthermore, when we degranulated these Slc37a2-knocked down BMMCs, no significant impairment in β-hexosaminidase release was seen compared to control siRNA treated BMMCs (Fig. 3d). Notably, 48-hours after degranulation Slc37a2-downregulated BMMCs had significantly impaired regranulation compared to controls (Fig. 3e). Furthermore, when we sought to visualize granules 48 hours after degranulation in siRNA treated BMMCs using an antibody to the granule component serotonin as a probe, we found that, in contrast to control BMMCs, very few serotonin granules were observed in Slc37a2 down regulated BMMCs (Fig. 3f). Taken together, these findings reveal that Slc37a2 is necessary for MC regranulation.

### Slc37a2 concentrates extracellular medium contents acquired by fluid phase endocytosis in endosomes

To deduce the role of Slc37a2 in MC regranulation, we first sought to visualize Slc37a2 localization both before and in the early stages of regranulation employing microscopy. For these studies, we opted to utilize mouse peritoneal MCs which are markedly more granulated than cultured BMMCs^42^. We sought to determine the localization of Slc37a2 in unstimulated, naïve MCs and recently degranulated MCs in relation to their granules using antibodies to CD63to mark MC granule membranes, and avidin-FITC, to mark MC granular components. Remarkably, in naïve MCs, Slc37a2 appeared to localize within MC granules that were encased in CD63^+^ membranes (Fig. 4a, top row) indicating that it was a regular MC granule component. As expected, 1 hour post degranulation, MCs appeared fully degranulated as no Avidin-FITC staining residual granules were detected (Fig. 4a, middle row). Consistent with this finding, most of the CD63^+^ membranes appeared to be localized at the cell periphery^43^. These cells also appeared to stain poorly for Slc37a2, indicating that the cellular content of Slc37a2 was markedly reduced with the exteriorization of MC granules. However, at 6 hours post-degranulation, which corresponded to the peak of Slc37a2 expression (Fig. 3b), newly formed Slc37a2 appeared localized to the periphery of the cell. Notably, this protein was not associated with CD63^+^ granule membranes (Fig. 4a, bottom row) suggesting that it had a function independent of the mature MC granule. In a previous study, when dextran was added to cell medium where Slc37a2 was over expressed in COS cells, dextran become specifically concentrated within Slc37a2^+^ vesicles, suggesting that Slc37a2 has the capacity to concentrate extracellular material taken up by fluid phase endocytosis and concentrate them within endosomes^44^. To see if Slc37a2 plays a similar role in MCs, we stably over expressed Slc37a2-EGFP in an immortalized rat MC line, RBL-2H3. We incubated 10,000 MW Texas Red dextran with Slc37a2-EGFP expressing RBL-2H3 and found that dextran was concentrated within distinct Slc37a2^+^ endosomes in MCs (Fig. 4b). To extend these findings, we knocked down expression of Slc37a2 in BMMCs using siRNAs then incubated these BMMCs with Texas Red dextran overnight. We found that in Slc37a2 knocked down BMMCs, there was significantly less internalized dextran compared to control siRNA treated cells (Fig. 4c). These data reveal that following MC degranulation, recently produced Slc37a2 is capable of concentrating extracellular dextran and potentially other extracellular nutrients in endosomes.

**Figure 4.**
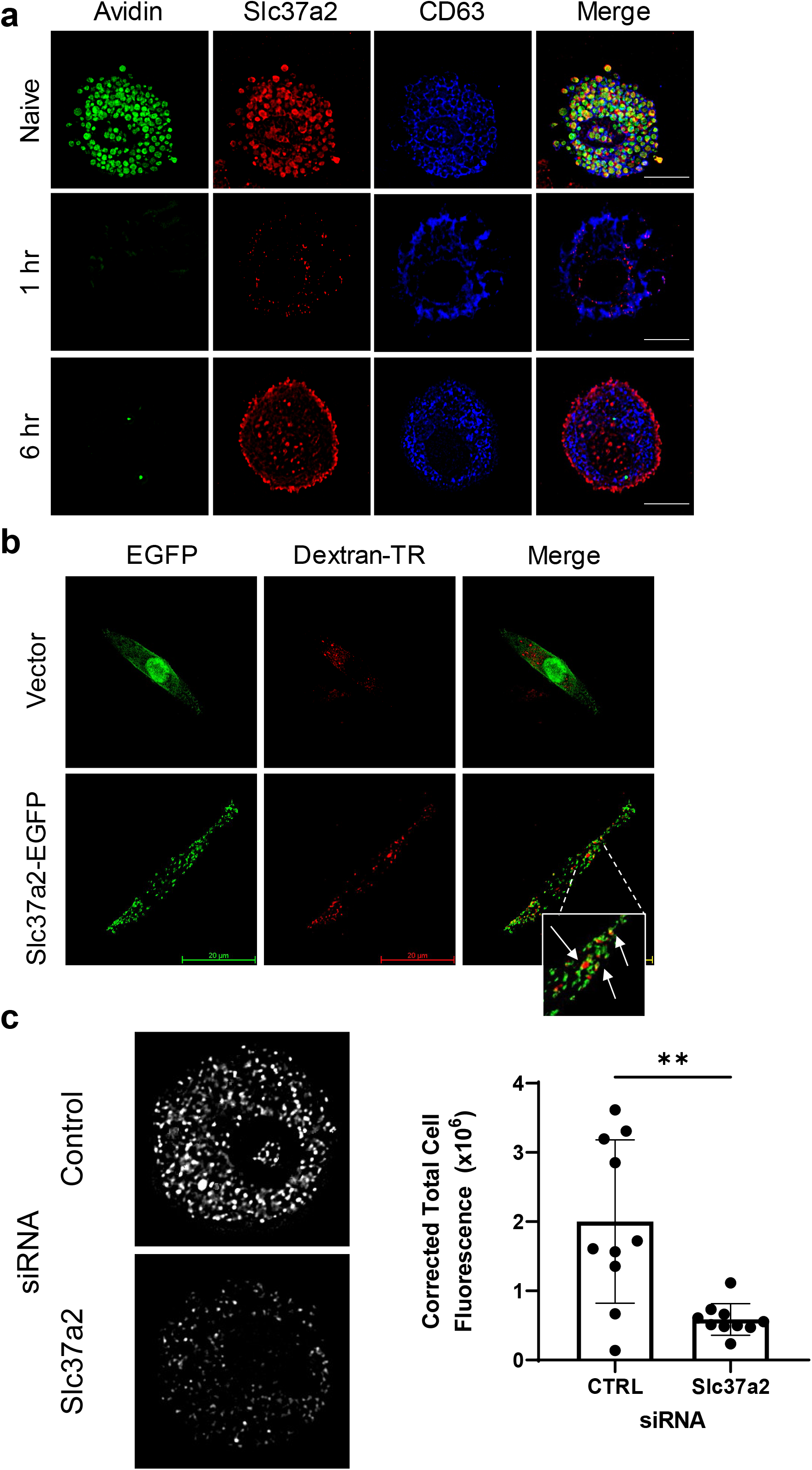
Slc37a2 localizes to endosomes during MC regranulation where it concentrates dextran. (**a**) Peritoneal lavages from C57BL/6J mice were obtained and treated with 1 μg/mL ionomycin for 1 hr in Tyrode’s buffer, then washed three times and resuspended in cell medium. Cells were cytospun onto coverslips at 0 (Naive), 1 hr, and 6 hrs post activation with ionomycin followed by fixation and staining for Avidin (green), Slc37a2 (red), and CD63 (blue). Scale bars: 10 μm. (**b**) RBLs were transduced with lentivirus expressing EGFP or Slc37a2-EGFP. Cells were plated onto coverslip and incubated overnight with 0.5 mg/mL dextran Texas Red 10,000 MW in cell medium. Cells were then washed three times and incubated in cell medium for 2 hrs, then fixed and visualized by microscopy. Arrows indicate Texas Red Dextran found within Slc37a2^+^ vesicles. All Images are representative of at least 2 independent experiments. (**c**) BMMCs were treated with Slc37a2 siRNAs or control siRNAs. 48 hrs after treatment, cells were incubated overnight with 0.5 mg/mL dextran Texas Red 10,000 MW in cell medium. Cells were then washed three times and incubated in cell medium for 2 hrs. Cells were then cytospun onto coverslips, then fixed and visualized by microscopy. Texas Red dextran was quantified using ImageJ; (*n* = 10). Data were analyzed using a Student’s *t* test. \*\**P* < 0.01.

### Peripherally located Slc37a2 associates with and activates mTOR

Since both mTORC1 and Slc37a2 are essential for regranulation, we wondered if these MC components interact with each other. In view of our finding that Slc37a2 concentrates extracellular nutrients within endosomes, it is conceivable that this activity may result in mTORC1 activation, analogous to how late endosomes signal to mTORC1^45^. Support for this notion comes from our finding that at 6 hours post degranulation, a subset of mTOR colocalized with nutrient bearing Slc37a2^+^ vesicles at the periphery of the MC (Fig. 5a). To see if Slc37a2 contributes to mTOR activation, we compared phosphorylation of mTOR following degranulation in control siRNA treated BMMCs and Slc37a2 knocked down BMMCs. While control siRNA treated BMMCs displayed increased mTOR phosphorylation by 2 hours post degranulation, Slc37a2 knocked down BMMCs exhibited limited mTOR phosphorylation (Fig. 5b), suggesting mTOR phosphorylation requires Slc37a2.

**Figure 5.**
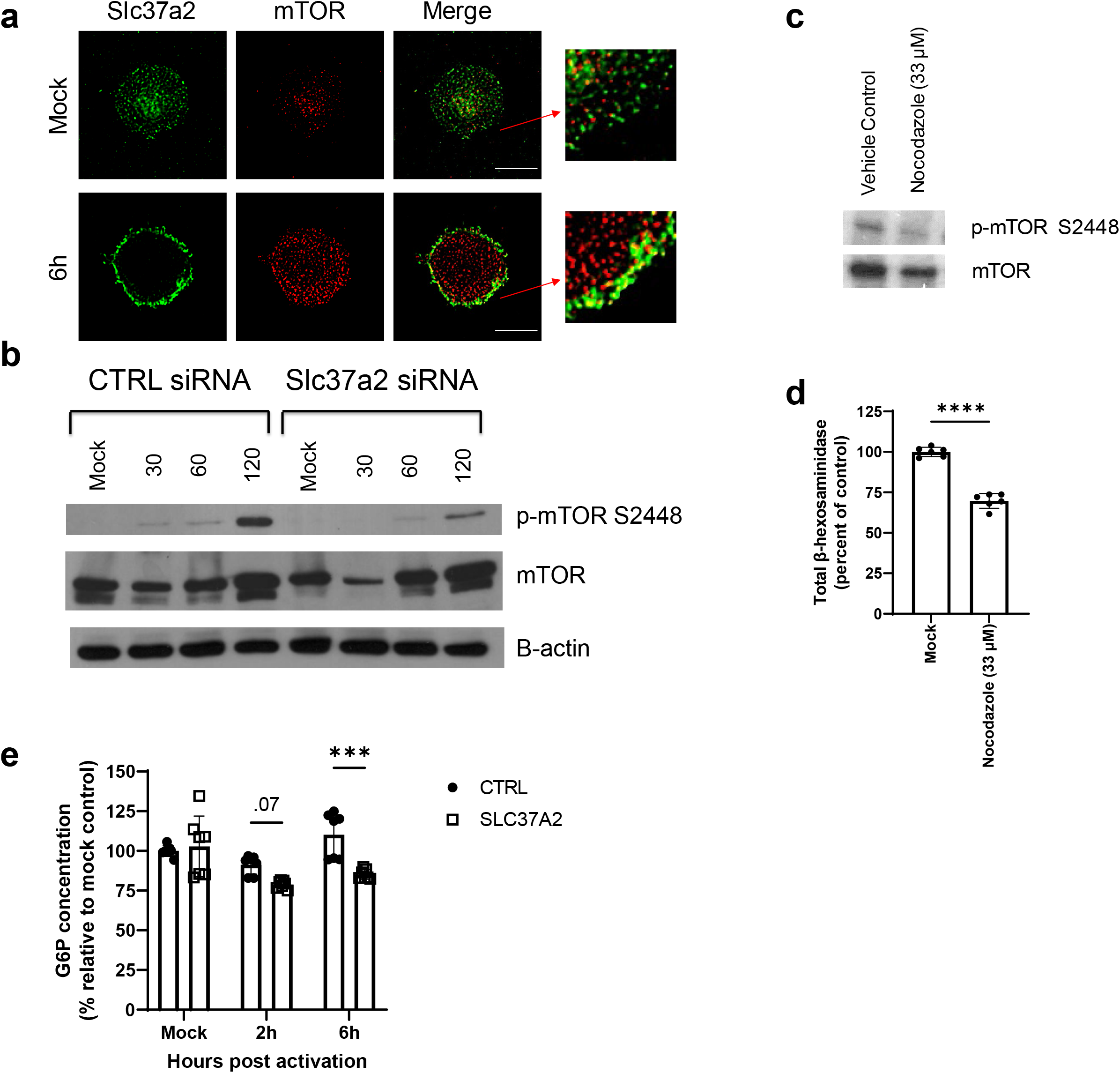
Trafficking of Slc37a2 to endosomes is necessary for mTOR activation. (**a**) BMMCs were activated using 1 μg/mL ionomycin in Tyrode’s buffer or Tyrode’s buffer alone (Mock) for 1 hr, then cells were washed three times and resuspended in cell medium. Six hrs post activation, BMMCs were cytospun onto coverslips fixed and stained with anti-Slc37a2 (green) and anti-mTOR (red). (**b**) BMMCs were treated with Slc37a2 siRNAs or control siRNAs. 48 hrs after treatment, cells were activated using 1 μg/mL ionomycin in Tyrode’s buffer or Tyrode’s buffer alone (Mock) for 1 hr then washed three times and resuspended in cell medium. Cells were lysed at 0 (Mock), 30, 60, and 120 min after activation and analyzed by Western blot. Immunoblots were incubated overnight with anti-p-mTOR S2448, anti-mTOR and anti-β-actin. Two independent experiments were performed with similar results. (**c-d**) BMMCs were activated using 1 μg/mL ionomycin in Tyrode’s buffer for 1 hr then washed three times and resuspended in cell medium containing 33 μM nocodazole or DMSO (vehicle control). (**c**) 120 min after activation, cells were lysed and analyzed by Western blot. Immunoblots were incubated overnight with anti-p-mTOR S2448 and anti-mTOR. (**d**) Total β-hexosaminidase was measured 48 hrs later. Results were normalized to vehicle control. Data are representative of 3 independent experiments. (**e**) BMMCs were treated with Slc37a2 siRNAs or control siRNAs. 48 hrs after treatment, cells were activated using 1 μg/mL ionomycin in Tyrode’s buffer for 1 hr then washed three times and resuspended in cell medium. G6P concentrations were determined in cell lysates at 0 (Mock), 2 hrs and 6 hrs post activation by colorimetric assay. Results were normalized to untreated control siRNA BMMCs. Data were analyzed using a Student’s *t* test (**d**) or a 2-way ANOVA with a Bonferroni post-test (**e**) \*\*\**P* < 0.001, \*\*\*\**P* < 0.0001.

To further support the notion that co-localization of Slc37a2 with mTOR in the cell periphery is critical to mTOR phosphorylation and subsequent MC regranulation, we sought to block the trafficking of newly synthesized Slc37a2 to the cell periphery by treating BMMCs after degranulation with nocodazole, a microtubule depolymerizer^46^. When BMMCs were treated with 33 μM nocodazole post degranulation, recently synthesized Slc37a2 accumulated in the perinuclear region of the cell (Supplemental Fig. 3a) whereas in vehicle control treated BMMCs Slc37a2 localized as expected to the cell periphery. In contrast to control treated BMMCs, mTOR in nocodazole treated BMMCs failed to get phosphorylated (Fig 5c) and these cells regranulated poorly (Fig. 5d), indicating that localization of Slc37a2 at the periphery of MCs during regranulation may be critical for mTOR phosphorylation and subsequent regranulation.

Previous studies have shown that G6P signals glucose levels to mTORC1 in cardiomyocytes^47,48^. In addition, Slc37a2 has the capacity to transport G6P^40^. Therefore, we wondered whether Slc37a2 increased intracellular G6P during regranulation to regulate mTOR activation in MCs. To test this, we measured intracellular G6P concentrations in Slc37a2 siRNA treated BMMCs before degranulation and at 2 hours and 6 hours post degranulation and compared them to control siRNA treated BMMCs. We found that there was no significant difference in intracellular G6P between Slc37a2 siRNA treated BMMCs compared to control siRNA treated cells prior to degranulation. Two hours post activation by ionomycin, Slc37a2 siRNA treated cells trended towards lower G6P concentrations (p=0.07) compared to control siRNA treated cells. Finally, at 6 hours post activation, there was a significantly lower concentration of G6P in Slc37a2 siRNA treated cells compared to controls (Fig. 5e). These results suggest that Slc37a2 is necessary to increase intracellular G6P during regranulation.

### Slc37a2 directs metabolic reprogramming linked to MC regranulation

The apparent capacity of Slc37a2 to concentrate nutrients from the extracellular medium brought into the cell via fluid phase endocytosis and the apparent capacity of the resulting endosomes to associate with mTOR, leading to mTOR phosphorylation, suggests that Slc37a2 may play a critical role in regulating MC energy levels to accommodate MC regranulation. Indeed, previous studies have shown that Slc37a2 regulates glycolysis in macrophages^41^ and the related protein G6PT (Slc37a4) has been shown to be necessary in maintaining G6P, lactate, and ATP amounts in neutrophils^49^. Since Slc37a2 has the capacity to transport G6P across membranes^40^, we first tested whether blocking G6P metabolism could inhibit regranulation. To do this, we used 2-deoxyglucose (2DG), a glucose analog that can enter cells and become phosphorylated into 2DG-6-phosphate, an analog of G6P but cannot be metabolized further. We found that treating BMMCs with 1 mM of 2DG 1 hour after degranulation significantly reduced its capacity to regranulate after degranulation (Fig. 6a) suggesting that glycolysis is indeed necessary for regranulation.

**Figure 6.**
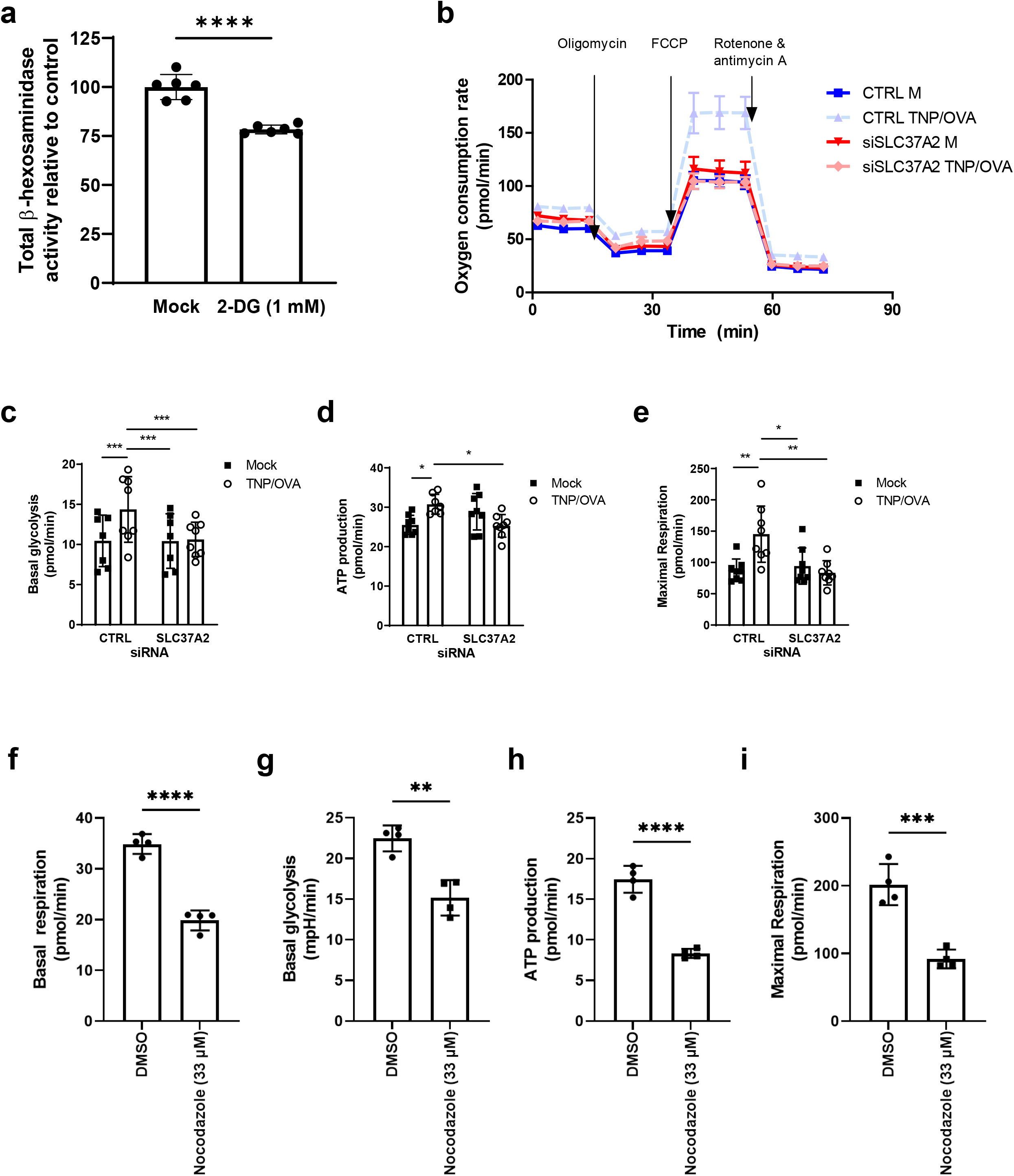
Slc37a2 trafficking to endosomes is necessary for metabolic reprogramming that occurs during MC regranulation. (**a**) BMMCs were activated using 1 μg/mL ionomycin in Tyrode’s buffer for 1 hr. Cells were then washed three times and resuspended in cell medium containing 1 mM 2-DG or cell medium alone (Mock). Total β-hexosaminidase was measured 48 hrs later. Data are representative of 2 independent experiments. (**b-e**) BMMCs were treated with Slc37a2 siRNAs or control siRNAs. 24 hrs after treatment, control and Slc37a2 siRNA treated cells were incubated with TNP-specific IgE in cell medium overnight. Cells were then washed three times and resuspended in Tyrode’s buffer containing TNP-OVA or cell medium alone (Mock) for 1 hr. Cells were then washed three times and resuspended in cell medium. (**b**) At 6 hrs post TNP-OVA treatment, cells were analyzed using the Seahorse XF using a mitochondrial stress test. (**c**) Basal glycolysis, (**d**) ATP production, and (**e**) maximal respiration were quantified. (**f-i**) BMMCs were activated using ionomycin in Tyrode’s buffer for 1 hr. Cells were then washed three times and resuspended in cell medium containing 33 μM nocodazole or DMSO (vehicle). 6 hrs post activation, cells were analyzed using the Seahorse XF using a mitochondrial stress test. (**f**) Basal respiration, (**g**) basal glycolysis, (**h**) ATP production, and (**i**) maximal respiration were quantified. Data were analyzed by a 1-way ANOVA with a Tukey’s post-test (**c-e**), or a Student’s *t* test (f-i). \**P* < 0.05; \*\**P* < 0.01; \*\*\**P* < 0.001; \*\*\*\**P* < 0.0001.

We next sought to investigate the role Slc37a2 has in glycolysis and oxidative phosphorylation during regranulation. To do this, we used Seahorse XF, which allows for measurements of extracellular acidification rate and oxygen consumption rate, relating to glycolysis and oxidative phosphorylation, respectively (Fig. 6b). In these experiments, we treated BMMCs with Slc37a2 siRNAs or control siRNAs and performed a mitochondrial stress test on these cells prior to degranulation or six hours after MC degranulation, corresponding to the peak expression of Slc37a2 in WT BMMCs. From these experiments we found that basal glycolysis is increased six hours post degranulation in control siRNA treated BMMCs, but a corresponding increase in Slc37a2 siRNA treated cells was not observed (Fig. 6c). Additionally, ATP production is increased six hours post degranulation in control siRNA treated cells, but not in Slc37a2 siRNA treated cells (Fig. 6d). Notably, maximum respiration is significantly increased in control siRNA treated BMMCs six hours post degranulation, but not in Slc37a2 siRNA treated BMMCs (Fig. 6e).

Additionally, to demonstrate the importance Slc37a2 localization on peripheral endosomes to metabolic reprogramming during regranulation, a mitochondrial stress test was performed 6 hours post degranulation on nocodazole treated BMMCs (Supplemental Fig. 3b). We found that both basal respiration and glycolysis were inhibited when Slc37a2 localization was blocked (Fig 6f-g). In addition, ATP production and maximum respiration were significantly reduced (Fig. 6h-i). These results suggest that the specific localization of Slc37a2 to peripheral endosomes during regranulation is critical for the metabolic reprogramming that occurs during regranulation. Furthermore, the presence of Slc37a2 in the cell periphery is necessary for increases in glycolysis, ATP production, and maximum respiration associated with the metabolic shift linked to MC regranulation.

### Slc37a2 conveys components of the extracellular medium into MC granules

Notably, in naïve primary MCs, Slc37a2 is found within mature MC granules (Fig 4a) suggesting that that the peripherally located Slc37a2 ultimately traffics to and become incorporated in nascent MC granules. To investigate if SLC37a2 associates with MC granule contents, we transduced the RBL-2H3 MC line expressing serglycin mCherry with plasmids expressing either Slc37a2-EGFP or an EGFP empty vector control. Serglycin is a proteoglycan found exclusively within the core of MC granules and in serglycin mCherry expressing RBL-2H3 cells, the granules can be readily visualized. Both sets of transduced serglycin mCherry RBL-2H3 cells were probed for CD63 and visualized by microscopy. As expected, serglycin was localized within CD63-marked vesicles in EGFP empty vector control transduced RBL-2H3 cells (Fig. 7a). Interestingly in Slc37a2 EGFP expressing RBL-2H3 cells, serglycin appeared encased in Slc37a2^+^ vesicles with CD63^+^ membranes seemingly serving to encase granules comprised of serglycin and Slc37a2^+^ vesicles. Presumably, Slc37a2^+^ vesicles appeared to have a stronger affinity for MC granules than CD63^+^-vesicles that typically encase MC granules (Fig. 7a). Thus, peripherally located Slc37a2^+^ vesicles may be destined to become components of MC granules.

**Figure 7.**
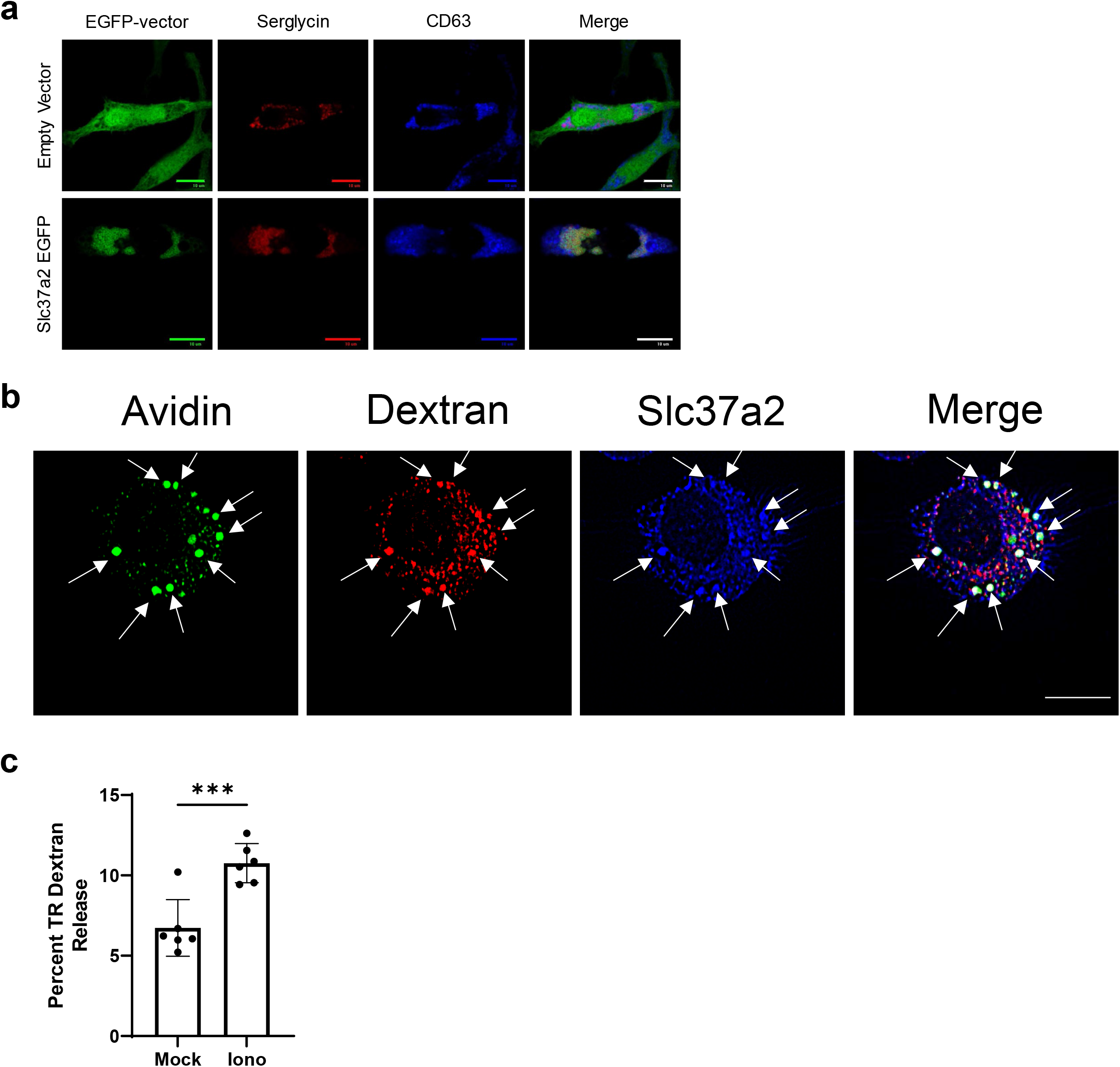
Slc37a2 traffics to nascent granules, carrying dextran into newly formed granules. (**a**) RBLs stably expressing serglycin mCherry (red) were transduced with lentivirus expressing EGFP or Slc37a2-EGFP (green). Cells were plated onto coverslip, fixed and stained with CD63 (blue). (**b**) BMMCs were activated using 1 μg/mL ionomycin in Tyrode’s buffer for 1 hr. Cells were then washed three times and resuspended in cell medium containing Texas Red Dextran (red). 48 hrs after activation, cells were cytospun onto coverslips, fixed, and stained with Avidin (green) and Slc37a2 (blue). Arrows indicate colocalization of Avidin, dextran, and Slc37a2. (**c**) BMMCs were incubated with Texas Red dextran in cell medium overnight. Cells were washed three times then treated with 1 μg/mL ionomycin for 1 hr in Tyrode’s buffer or Tyrode’s buffer alone (Mock). Percent released dextran was determined by quantifying fluorescence signal in supernatant and comparing it to total fluorescence in supernatant and lysed pellet. Results are representative of two independent experiments. Data were analyzed using a Student’s *t* test. \*\*\**P* < 0.001.

If SLC37a2 vesicles are trafficked into MC granules, then cargo acquired from the extracellular medium that are stored and concentrated in these vesicles may also be incorporated into MC granules. To investigate this possibility, we degranulated peritoneal MCs and exposed them to Texas Red dextran, a marker of nondescript extracellular medium components. We found that at 48 hrs, Slc37a2 was associated with nascent Avidin^+^ granules. Furthermore, these granules were also associated with dextran (Fig. 7b). This suggests that extracellular components are trafficked in Slc37a2^+^ endosomes into newly formed granules. To further confirm this, we incubated BMMCS with Texas Red dextran for 24 hours and then degranulated these BMMCs using ionomycin. We found that a significantly increased amount of dextran was released along with granules compared to unactivated BMMC controls (Fig. 7c).

Thus taken together, Slc37a2 appears to have an additional property of concentrating cargo acquired from the extracellular medium into vesicles that are directly trafficked into MC granules, suggesting that Slc37a2 has the capacity to regulate the composition of MC granules.

## Discussion

MCs are best known for their high content of electron-dense secretory granules that occupy most of the cytoplasm of the cell^10^. These granules are replete with a panoply of pharmacologically active agents, and the MC’s capacity to simultaneously exocytose most of their granules upon activation is one of the primary reasons that MCs are regarded as the early triggers of a wide range of inflammatory disorders such as asthma, arthritis, and anaphylaxis. Although MCs were shown to regranulate *in vitro* over 50 years ago^13^, this is the first indication of regranulation *in vivo*. Importantly, we found that following IgE mediated activation, MCs are capable of regranulating and repeatedly mediating anaphylaxis with inflammatory levels comparable to naïve MCs activated for the first time. This shows that the regranulation capacity of MCs *in vivo* is comprehensive and restores function. That MCs possess the innate capacity to undergo at least two rounds of regranulation and degranulation *in vivo* after an initial degranulation response would suggest that in addition to initiating inflammatory responses upon activation, MCs can contribute to sustaining these inflammatory activities for as long as the activating agent is present.

While MCs maintained their capacity to undergo multiple cycles of degranulation and regranulation *in vivo*, changes to the inflammatory response were observed in the second and third MC activation. Our studies were specifically designed to reveal IgE mediated MC degranulation, and to exclude the contribution of other immune cells, such as lymphocytes capable of memory responses. Rather than actively immunizing test mice with OVA Ag, we passively immunized them with OVA-specific IgE antibodies so that, for the most part, only MCs were sensitized and capable of reacting to each challenge with Ag. Furthermore, Ag challenges of the mice were undertaken 7 days apart by which time all of the inflammation induced by the previous challenge had subsided. Thus, the immune responses evoked after each challenge were primarily a result of MCs with little or no contribution from innate or adaptive immune cells. Notably, an influx of monocytes was observed after each Ag challenge, however, neutrophil recruitment was reduced in the second and third Ag challenges. This matches the trend towards an increased rate of recovery from anaphylaxis after each subsequent Ag challenge and suggests a possible adaptation of MCs to repeat Ag challenge. Interestingly, a similar observation was made after repeat FcεRI triggering in human cord blood MCs *in vitro* resulted in modulated MC function^50^.

This capacity of MCs to undergo multiple cycles of degranulation and regranulation is unique. While much is known regarding how MCs degranulate and release their granules, comparatively, little is known regarding the molecular events associated with initial MC granule formation, and no information is available on granule reformation. Conceivably, for MCs to fully regenerate the repertoire of granules populating their cytosol, significant protein and lipid synthesis in concert with intracellular vesicle trafficking and packaging of various granule components into the granule structure would need to occur. These activities require significant energy and so MCs would need to undergo significant metabolic reprogramming. Our studies have implicated the protein kinase mTOR, the well-known mediator of cellular metabolism, growth, and proliferation as the cellular switch initiating MC regranulation associated activities^51^. This conclusion is based on the findings that (i) mTOR and S6K a serine/threonine kinase, a signaling substrate downstream of mTOR, become phosphorylated following MC degranulation, (ii) impeding mTOR activation with a chemical inhibitor, torin, prevents MCs from regranulation *in vitro* without impeding their capacity to degranulate and (iii) genetically knocking out Raptor, a necessary gene for mTORC1 expression, blocks the capacity of MCs to regranulate *in vivo* without affecting their capacity to degranulate. Additionally, mTOR activation during MC regranulation was closely linked to increased glycolysis, ATP production and maximum respiration in MCs (Fig. 6). That mTOR is important for MC regranulation following MC degranulation is consistent with previous reports showing that activation of mTOR in MCs inhibited IgE mediated MC degranulation and cytokine release as well as their capacity to proliferate^52,53^. Presumably, activating mTOR triggers a metabolic switch in steady state MCs to accommodate regranulation which interferes with both degranulation and MC proliferation activities.

A critical signaling substrate involved in mTOR mediated MC regranulation is Slc37a2. This molecule was discovered when we employed an unbiased approach to uncover substrates relevant to MC regranulation. This approach involved examining MCs for metabolic genes that were highly upregulated when MC degranulation had subsided. Slc37a2, a transmembrane protein implicated previously in G6P transport in macrophages^41^ was the most upregulated gene involved in glucose metabolism that we encountered (Fig. 3a). Indeed, 6 hours after MC degranulation, Slc37a2 expression was found to have increased 400-fold over baseline (Fig. 3b). Furthermore, this newly produced Slc37a2 was found to specifically localize at the MC periphery and on vesicles that had recently endocytosed nutrients found in the extracellular medium (Fig. 4). MC uptake of labelled extracellular dextran is analogous to findings in Slc37a2 over-expressing COS cells where dextran from the extracellular medium was found to be harbored within Slc37a2+ endosomes^44^. Conceivably, peripherally located Slc37a2 contributes to the uptake of nutrients from the extracellular medium. This is supported by our studies showing that nocodazole treatment of BMMCs, which prevents Slc37a2 trafficking to endosomes, reduced basal respiration, basal glycolysis, ATP production, and maximal respiration 6 hours post degranulation, similar to what is observed when Slc37a2 is knocked down by siRNAs (Fig. 6 c-i).

So how are these extracellular nutrient-encapsulating Slc37a2+ endocytic compartments in MCs connected to mTOR mediated metabolic switch and MC regranulation? One possibility is that cargo encased in Slc37a2+ endosomes signals mTOR in a manner analogous to how amino acids, glucose, and ATP borne in lysosomes activates the cellular nutrient sensor mTOR found associating with the lysosome surface and triggering a metabolic switch^51^. Support for this possibility comes from the finding that nutrient-bearing Slc37a2+ endosomes found in the cell periphery appear to associate with mTOR. Furthermore, knockdown of Slc37a2 results in reduced G6P and ATP within MCs during regranulation, two critical signals for mTORC1 activation. Finally, knocking down of Slc37a2 or preventing the trafficking of Slc37a2 to the cell periphery where it makes contact with mTOR was found to prevent MC regranulation, emphasizing the critical nature of the Slc37a2-mTOR interactions to this activity.

Another intriguing finding was that during MC regranulation, peripherally located Slc37a2^+^ endosomes travel to nascent MC granules or progranules which are located in the “progranule zone” to become MC granule components. While we are not able to ascertain its function within the granule, one possibility is that Slc37a2 contributes to concentrating granule components making it more compact and particulate, a phenomenon sometimes known as condensation^19^. This role for SLC37a2 was recently observed when intracellular Slc37a2^+^ vesicles within microglia were found to exhibit the capacity to concentrate vesicular contents and reduce vesicle volume to a fraction of its original size. Loss of Slc37a2 blocked this vesicular shrinkage, resulting in its expansion and bloating of the cell^54^.

What drives SLC37a2^+^ endosomes to traffic into nascent MC granules during regranulation is unclear. When SLC37a2 was over expressed in RBL-2H3 cells, SLC37a2 rapidly gathered around MC granules and seemed to displace the CD63^+^ membranes from around granule cores (Fig. 7a). Apparently, SLC37a2 has a stronger affinity for the serglycin-rich and negatively charged granule core than CD63^+^ membranes. It is noteworthy that when SLC37a2^+^ endosomes fuse with nascent MC granules they do so bearing cargo endocytosed from the extracellular medium. Indeed, dextran acquired from the extracellular medium by SLC37a2^+^ vesicles were translocated to nascent MC granules within 48 hours of degranulation. That this dextran was incorporated into MC granules was confirmed by the observation that when MCs were activated, dextran was released along with granules. This finding is consistent with previous claims that histamine and TNF found in the extracellular medium could be incorporated into MC granules and released when MCs degranulate^55,56^, but the underlying mechanism has remained a mystery until now. These studies also reveal that Slc37a2 is potentially a key regulator of MC granule composition. Previous studies have reported that mature MC granules are formed when progranules leave the “progranule zone” near the golgi and endoplasmic reticulum and fuse with other progranules, immature granules, endosomes, or lysosomes. Our studies reveal for the first time that regranulation also involve the active participation of vesicles recruited from the plasma membrane bearing cargo from the extracellular medium and reveal the critical role of Slc37a2 as a regulator of MC granule content.

Finally, now that we have demonstrated that MCs have the capacity to regranulate *in vivo*, future studies can be designed to assess the specific contribution of the regranulation property to the pathogenesis of MCs in various inflammatory disorders. If this property is deemed to be significant, selectively targeting it may be an efficient way to temper the excessive activities of MCs without impairing its beneficial role in promoting innate and adaptive immune responses to infectious agents and in maintaining homeostasis.

## Acknowledgements

We thank the Duke Cellular Metabolism Analysis Core, especially Amanda Nichols, for their expertise and support on our Seahorse XF experiments. We thank the Duke Microarray Core facility (a Duke NCI Cancer Institute and a Duke Genomic and Computational Biology shared resource facility) for their technical support, microarray data management and feedback on the generation of the microarray data reported in this manuscript. We would also like to thank the Duke Genomic Analysis and Bioinformatics Core and Dr. David Corcoran for their assistance with analyzing our microarray data.

## Competing Interests

The authors declare no competing interests.

## Notes

### Competing Interest Statement

The authors have declared no competing interest.

## References

1 Gurish, M. F. & Austen, K. F. Developmental origin and functional specialization of mast cell subsets. Immunity. (2012) 37, 25–33.

2 Metcalfe, D. D., Baram, D. & Mekori, Y. A. Mast cells. Physiological reviews. (1997) 77, 1033–1079.

3 Marshall, J. S. Mast-cell responses to pathogens. Nat Rev Immunol. (2004) 4, 787–799.

4 Abraham, S. N. & St John, A. L. Mast cell-orchestrated immunity to pathogens. Nat Rev Immunol. (2010) 10, 440–452.

5 Urb, M. & Sheppard, D. C. The role of mast cells in the defence against pathogens. PLoS Pathog. (2012) 8, e1002619.

6 Theoharides, T. C. & Kalogeromitros, D. The critical role of mast cells in allergy and inflammation. Ann N Y Acad Sci. (2006) 1088, 78–99.

7 Amin, K. The role of mast cells in allergic inflammation. Respir Med. (2012) 106, 9–14.

8 Boyce, J. A. The role of mast cells in asthma. Prostaglandins Leukot Essent Fatty Acids. (2003) 69, 195–205.

9 Yu, M. et al. Mast cells can promote the development of multiple features of chronic asthma in mice. J Clin Invest. (2006) 116, 1633–1641.

10 Wernersson, S. & Pejler, G. Mast cell secretory granules: armed for battle. Nat Rev Immunol. (2014) 14, 478–494.

11 Kiernan, J. A. Production and life span of cutaneous mast cells in young rats. J Anat. (1979) 128, 225–238.

12 Padawer, J. Mast cells: extended lifespan and lack of granule turnover under normal in vivo conditions. Exp Mol Pathol. (1974) 20, 269–280.

13 Kobayasi, T. & Asboe-Hansen, G. Degranulation and regranulation of human mast cells. An electron microscopic study of the whealing reaction in urticaria pigmentosa. Acta dermato-venereologica. (1969) 49, 369–381.

14 Xiang, Z., Block, M., Lofman, C. & Nilsson, G. IgE-mediated mast cell degranulation and recovery monitored by time-lapse photography. J Allergy Clin Immunol. (2001) 108, 116–121.

15 Ando, T. et al. Mast cells are required for full expression of allergen/SEB-induced skin inflammation. J Invest Dermatol. (2013) 133, 2695–2705.

16 Okayama, Y., Ra, C. & Saito, H. Role of mast cells in airway remodeling. Curr Opin Immunol. (2007) 19, 687–693.

17 Hammel, I. et al. Differences in the volume distributions of human lung mast cell granules and lipid bodies: evidence that the size of these organelles is regulated by distinct mechanisms. J Cell Biol. (1985) 100, 1488–1492.

18 Combs, J. W. Maturation of rat mast cells. An electron microscope study. J Cell Biol. (1966) 31, 563–575.

19 Hammel, I., Lagunoff, D. & Galli, S. J. Regulation of secretory granule size by the precise generation and fusion of unit granules. J Cell Mol Med. (2010) 14, 1904–1916.

20 Moon, T. C., Befus, A. D. & Kulka, M. Mast Cell Mediators: Their Differential Release and the Secretory Pathways Involved. Frontiers in Immunology. (2014) 5, 569.

21 Azouz, N. P. et al. Rab5 is a novel regulator of mast cell secretory granules: impact on size, cargo, and exocytosis. J Immunol. (2014) 192, 4043–4053.

22 Rönnberg, E., Melo, F. R. & Pejler, G. Mast cell proteoglycans. J Histochem Cytochem. (2012) 60, 950–962.

23 Kunder, C. A. et al. Mast cell-derived particles deliver peripheral signals to remote lymph nodes. J Exp Med. (2009) 206, 2455–2467.

24 St John, A. L., Chan, C. Y., Staats, H. F., Leong, K. W. & Abraham, S. N. Synthetic mast-cell granules as adjuvants to promote and polarize immunity in lymph nodes. Nat Mater. (2012) 11, 250–257.

25 Chi, H. Regulation and function of mTOR signalling in T cell fate decisions. Nature Reviews Immunology. (2012) 12, 325–338.

26 Salmond, R. J. mTOR Regulation of Glycolytic Metabolism in T Cells. Frontiers in Cell and Developmental Biology. (2018) 6.

27 Saxton, R. A. & Sabatini, D. M. mTOR Signaling in Growth, Metabolism, and Disease. Cell. (2017) 169, 361–371.

28 Finlay, D. Regulation of glucose metabolism in T cells: new insight into the role of Phosphoinositide 3-kinases. Frontiers in Immunology. (2012) 3.

29 Ang, W. X. et al. Mast cell desensitization inhibits calcium flux and aberrantly remodels actin. J Clin Invest. (2016) 126, 4103–4118.

30 Carvalho, B. S. & Irizarry, R. A. A framework for oligonucleotide microarray preprocessing. Bioinformatics. (2010) 26, 2363–2367.

31 Gentleman, R. C. et al. Bioconductor: open software development for computational biology and bioinformatics. Genome Biol. (2004) 5, R80.

32 Ritchie, M. E. et al. limma powers differential expression analyses for RNA-sequencing and microarray studies. Nucleic acids research. (2015) 43, e47.

33 Mootha, V. K. et al. PGC-1alpha-responsive genes involved in oxidative phosphorylation are coordinately downregulated in human diabetes. Nat Genet. (2003) 34, 267–273.

34 Korotkevich, G. et al. Fast gene set enrichment analysis. bioRxiv. (2021), 060012.

35 Choi, H. W. et al. Perivascular dendritic cells elicit anaphylaxis by relaying allergens to mast cells via microvesicles. Science. (2018) 362.

36 Tharp, M. D., Seelig, L. L., Jr., Tigelaar, R. E. & Bergstresser, P. R. Conjugated avidin binds to mast cell granules. J Histochem Cytochem. (1985) 33, 27–32.

37 Weichhart, T., Hengstschlager, M. & Linke, M. Regulation of innate immune cell function by mTOR. Nat Rev Immunol. (2015) 15, 599–614.

38 Liu, Q. et al. Development of ATP-competitive mTOR inhibitors. Methods Mol Biol. (2012) 821, 447–460.

39 Cappello, A. R., Curcio, R., Lappano, R., Maggiolini, M. & Dolce, V. The Physiopathological Role of the Exchangers Belonging to the SLC37 Family. Frontiers in chemistry. (2018) 6.

40 Pan, C. J. et al. SLC37A1 and SLC37A2 are phosphate-linked, glucose-6-phosphate antiporters. PloS one. (2011) 6, e23157.

41 Wang, Z. et al. Solute Carrier Family 37 Member 2 (SLC37A2) Negatively Regulates Murine Macrophage Inflammation by Controlling Glycolysis. iScience. (2020) 23, 101125.

42 Malbec, O. et al. Peritoneal Cell-Derived Mast Cells: An In Vitro Model of Mature Serosal-Type Mouse Mast Cells. The Journal of Immunology. (2007) 178, 6465–6475.

43 Schäfer, T., Starkl, P., Allard, C., Wolf, R. M. & Schweighoffer, T. A granular variant of CD63 is a regulator of repeated human mast cell degranulation. Allergy. (2010) 65, 1242–1255.

44 Kim, J. Y., Tillison, K., Zhou, S., Wu, Y. & Smas, C. M. The major facilitator superfamily member Slc37a2 is a novel macrophage- specific gene selectively expressed in obese white adipose tissue. American journal of physiology. Endocrinology and metabolism. (2007) 293, E110–120.

45 Flinn, R. J., Yan, Y., Goswami, S., Parker, P. J. & Backer, J. M. The late endosome is essential for mTORC1 signaling. Molecular biology of the cell. (2010) 21, 833–841.

46 Vasquez, R. J., Howell, B., Yvon, A. M., Wadsworth, P. & Cassimeris, L. Nanomolar concentrations of nocodazole alter microtubule dynamic instability in vivo and in vitro. Molecular biology of the cell. (1997) 8, 973–985.

47 Karlstaedt, A., Khanna, R., Thangam, M. & Taegtmeyer, H. Glucose 6-Phosphate Accumulates via Phosphoglucose Isomerase Inhibition in Heart Muscle. Circ Res. (2020) 126, 60–74.

48 Roberts, D. J., Tan-Sah, V. P., Ding, E. Y., Smith, J. M. & Miyamoto, S. Hexokinase-II positively regulates glucose starvation-induced autophagy through TORC1 inhibition. Mol Cell. (2014) 53, 521–533.

49 Jun, H. S., Weinstein, D. A., Lee, Y. M., Mansfield, B. C. & Chou, J. Y. Molecular mechanisms of neutrophil dysfunction in glycogen storage disease type Ib. Blood. (2014) 123, 2843–2853.

50 Suurmond, J. et al. Repeated FcepsilonRI triggering reveals modified mast cell function related to chronic allergic responses in tissue. J Allergy Clin Immunol. (2016) 138, 869–880.

51 Saxton, R. A. & Sabatini, D. M. mTOR Signaling in Growth, Metabolism, and Disease. Cell. (2017) 168, 960–976.

52 Blatt, K. et al. The PI3-Kinase/mTOR-Targeting Drug NVP-BEZ235 Inhibits Growth and IgE-Dependent Activation of Human Mast Cells and Basophils. PloS one. (2012) 7, e29925.

53 Rakhmanova, V., Jin, M. & Shin, J. Inhibition of Mast Cell Function and Proliferation by mTOR Activator MHY1485. Immune Netw. (2018) 18, e18.

54 Villani, A. et al. Clearance by Microglia Depends on Packaging of Phagosomes into a Unique Cellular Compartment. Developmental cell. (2019) 49, 77–88.e77.

55 Olszewski, M. B., Groot, A. J., Dastych, J. & Knol, E. F. TNF trafficking to human mast cell granules: mature chain-dependent endocytosis. J Immunol. (2007) 178, 5701–5709.

56 Ohtsu, H. et al. Plasma extravasation induced by dietary supplemented histamine in histamine-free mice. Eur J Immunol. (2002) 32, 1698–1708.

