## Supplemental Material for "Mast cell regranulation involves a metabolic switch promoted by the interaction between mTORC1 and a glucose-6-phosphate transporter"

Supplemental Figure 1

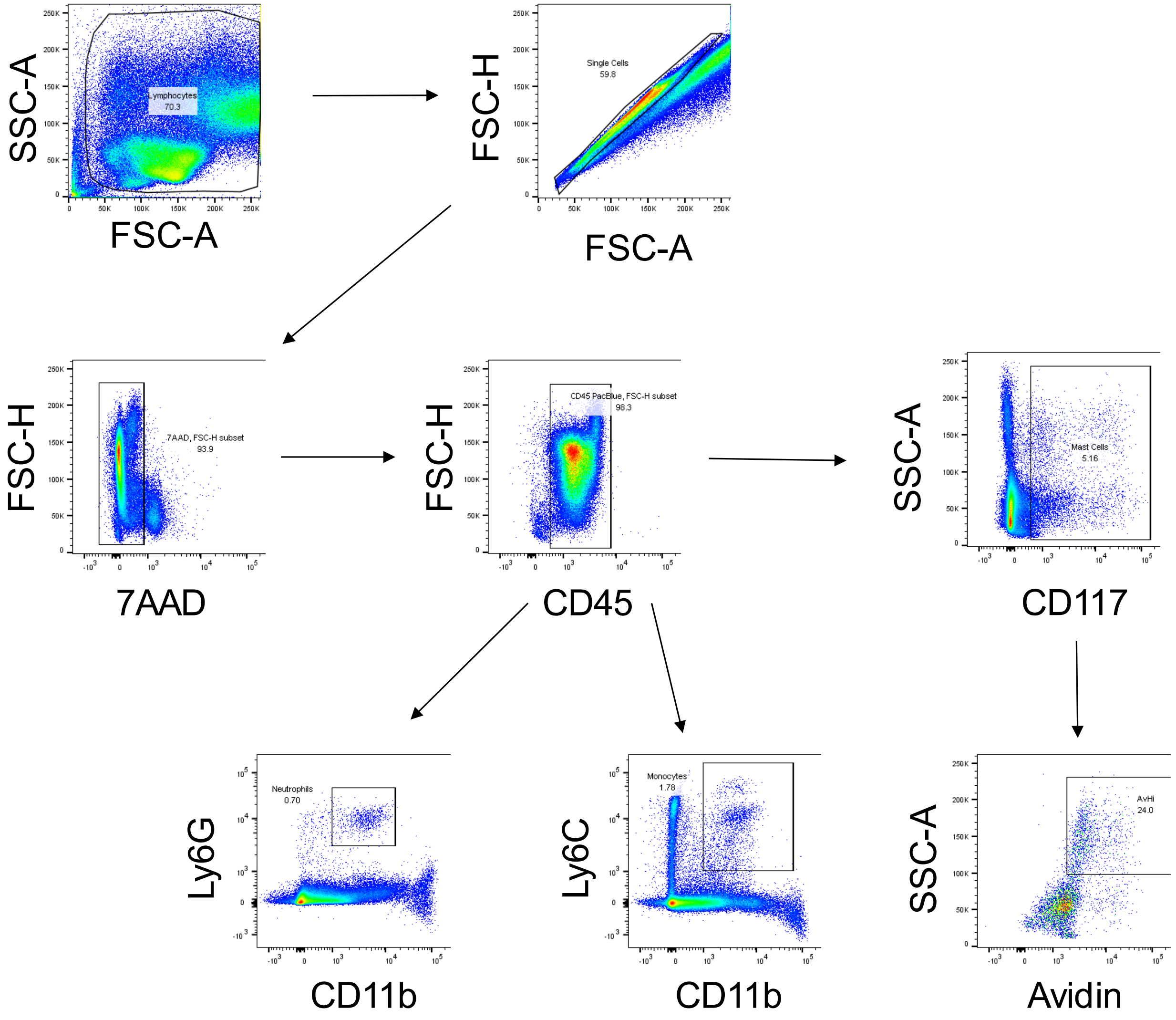


**Supplemental Figure 1. Gating Strategy for flow cytometry analysis of peritoneal lavages.**

Supplemental Figure 2

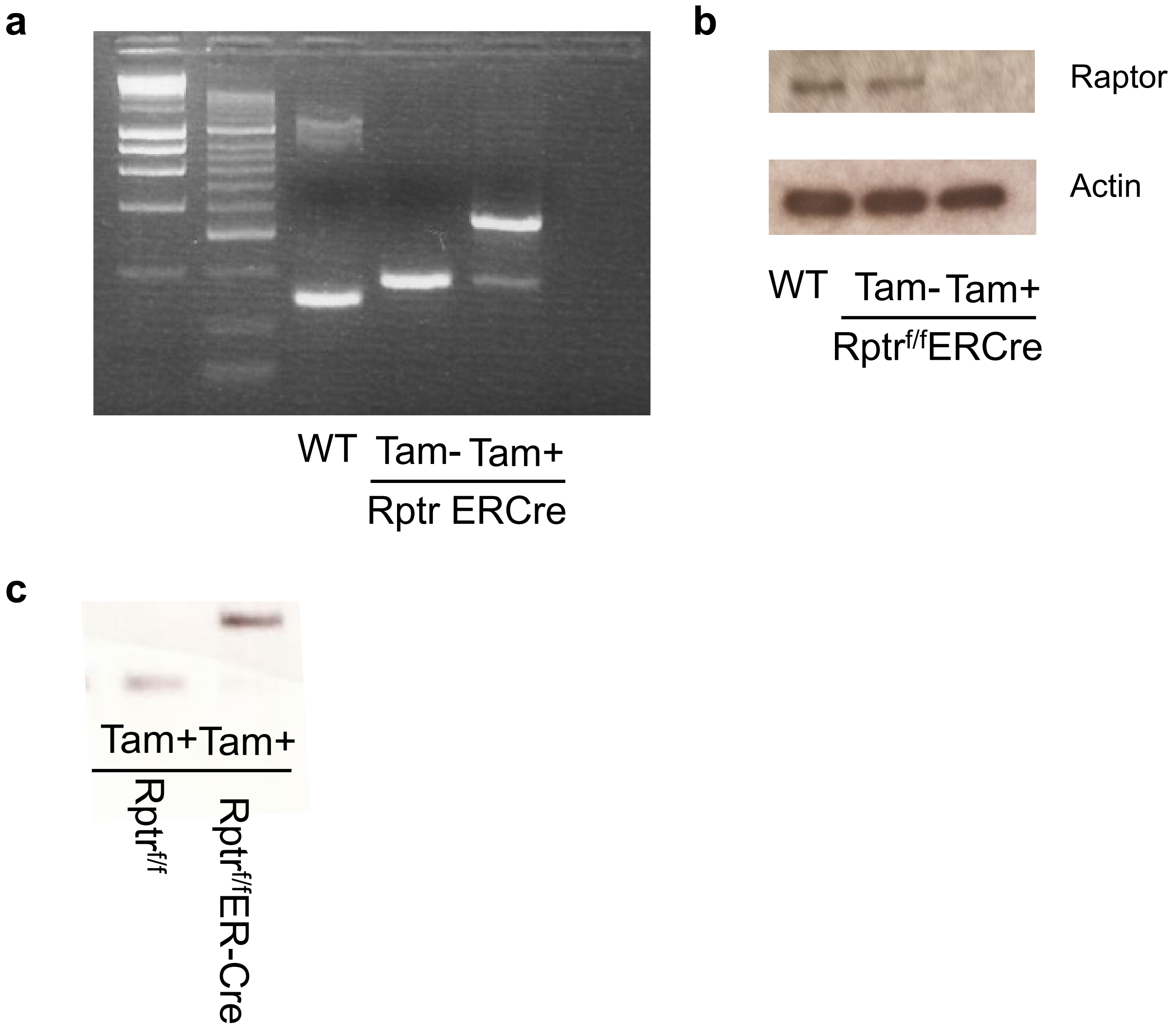


**Supplemental Figure 2. Tamoxifen treatment of Raptor ER-Cre.** (**a**) Raptor-T-KO mice were treated daily for 5 days with 2 µg in 100 µl PBS i.p. On day 7, blood from Raptor-T-KO mice, C57BL/6J (WT), and Raptor$^{f/f}$ ER-Cre mice untreated with tamoxifen were collected. DNA was extracted from samples and PCR amplified using primers surrounding exon 6 of Raptor. Red arrow indicates amplicon from Raptor knockout. Black arrow indicates intact floxed Raptor exon 6. (**b**) BMMCs from Raptor$^{f/f}$ or Raptor-T-KO mice were treated with 4-hydroxytamoxifen overnight. DNA was extracted and PCR amplified using primers surrounding exon 6 of Raptor. Red arrow indicates amplicon from Raptor knockout. Black arrow indicates intact floxed Raptor exon 6. (**c**) Western blot analysis of BMMCs from C5BL/6J (WT), Raptor$^{f/f}$ and Raptor-T-KO mice after 4-hydroxytamoxifen treatment. Immunoblots were incubated overnight with anti-Raptor or anti-β-actin.

### Supplemental Figure 3

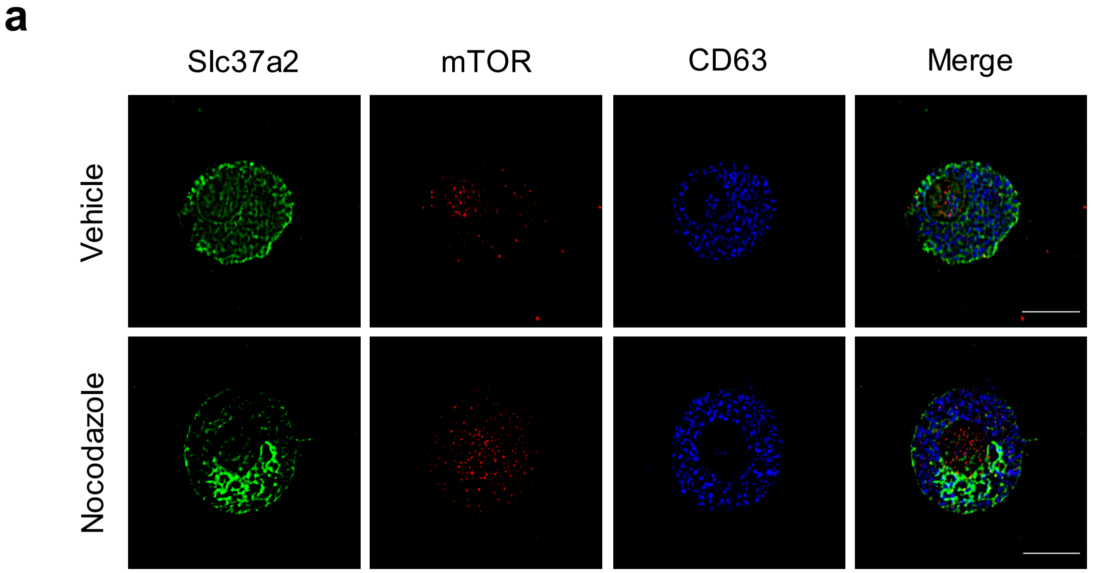


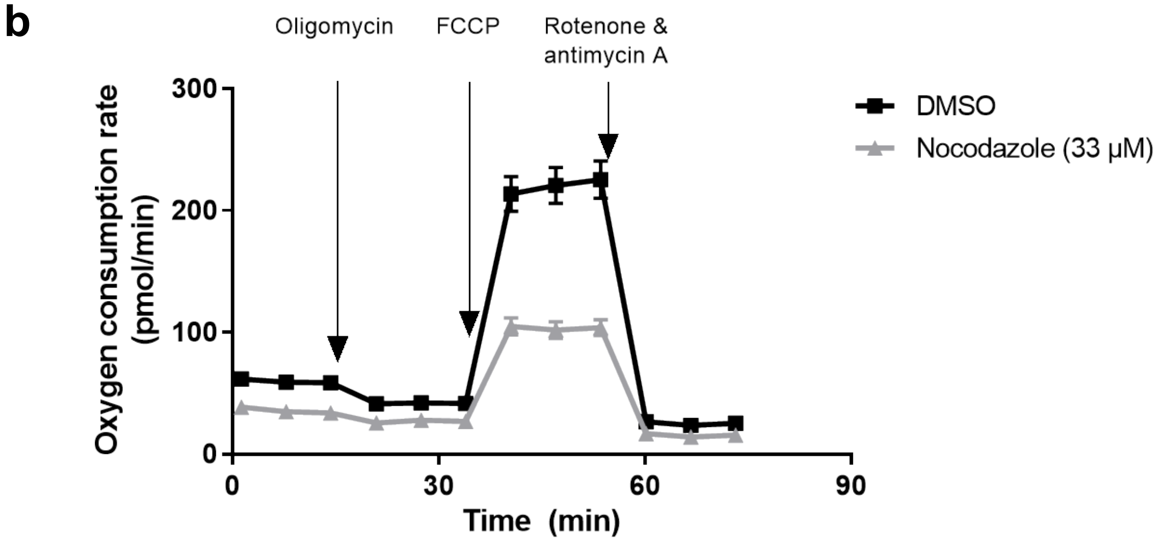


**Supplemental Figure 3. Nocodazole treatment of BMMCs.** (**a-b**) BMMCs were activated using 1 μg/mL ionomycin in Tyrode's buffer for 1 hr. Cells were washed three times and resuspended in cell medium containing nocodazole or DMSO (vehicle control). (**a**) Cells were cytospun onto slides 6 hrs post activation, then fixed and stained using Slc37a2 (green), mTOR (red), and CD63 (blue). (**b**) At 6 hrs post activation, cells were analyzed using a mitochondrial stress test on the Seahorse XF.
